# ALS/FTD-associated TDP-43 mutations promote fragility of genes governing excitatory neurotransmission via topoisomerase IIβ impairment

**DOI:** 10.64898/2025.12.08.693088

**Authors:** Harman Sharma, Sushma Koirala, Megan Fowler, Mary-Louise Rogers, Yee Lian Chew, Anna Konopka

## Abstract

Abnormal TAR DNA/RNA-binding protein 43 (TDP-43) is a hallmark of amyotrophic lateral sclerosis (ALS) and frontotemporal dementia (FTD), characterized by cytoplasmic mis-localization, aggregation, and pathogenic mutations. Altered excitatory neuronal transmission is an early functional defect in these diseases; however, the underlying mechanisms remain unclear. Neuronal activity can induce DNA double-strand breaks (DSBs), suggesting a potential link between altered excitability and genomic instability. Here, we demonstrate for the first time that TDP-43 is involved in neuronal activity-induced DSBs. Moreover, we show that ALS/FTD-associated TDP-43 mutations (A90V and A315T) disrupt this mechanism, leading to the accumulation of DSBs and altered neuronal activity compared to wild-type TDP-43 protein. Using the BLISS technique, we mapped genome-wide DSBs in primary mouse neurons expressing wild-type or mutant (A90V, A315T) TDP-43 and found enrichment of DSBs within genes regulating excitatory transmission. Mechanistically, the TDP-43 A90V mutation impairs topoisomerase IIβ function, resulting in enzyme trapping and/or DNA supercoiling that predisposes DNA to breaks. Notably, this impairment is partially rescued by tyrosyl-DNA phosphodiesterase 2 (Tdp2) overexpression. These findings uncover a novel mechanism linking aberrant neuronal activity and DNA damage, bridging two key pathological hallmarks of ALS/FTD associated with TDP-43 dysfunction. It also paves the way for developing novel therapeutic approaches that rely on targeted DSB repair and/or modulating topoisomerase IIβ-DNA complexes.

**Highlights:**

- ALS/FTD linked TDP-43 mutants, unlike wild type, induce DSBs in key neuronal genes
- TDP-43 driven DSBs impair neuronal transmission, linking two ALS/FTD pathologies
- TDP-43 pathology impairs topoisomerase IIβ, leading to DSB

## Background

TAR RNA/DNA binding protein 43 (TDP-43) is strongly associated with several neurodegenerative diseases (1). TDP-43 forms pathological aggregates in the cytoplasm of affected neurons in almost all cases of amyotrophic lateral sclerosis (ALS) (approximately 97%) (2), 50% of cases with frontotemporal lobe dementia (FTLD) (3) and 20-50% of cases with Alzheimer’s disease (AD) (4). In addition, the presence of TDP-43 mutations has been reported in both ALS and FTD (5).

Abnormal forms of TDP-43 have been linked to both altered neuronal excitability and accumulation of DNA damage in ALS (6–10); however, these phenomena have been considered independent rather than parts of a common mechanism. While the dysfunction of TDP-43 is enough to trigger neuronal hyperexcitability in mouse model of ALS (8), abnormal TDP-43 impairs the repair of the most lethal form of DNA damage, double stranded breaks (DSBs), causing genomic instability (6, 7, 11). Interestingly, emerging evidence links DSBs to neuronal excitability (12–14). Neuronal stimulation induces the formation of DSBs at the promoters of a subset of early-response genes, facilitating the regulation of their expression (12). Furthermore, a novel activity-dependent DNA repair mechanism has been identified in neurons that relies on the NuA4-TIP60 complex, linking synaptic activity to DNA repair (13). The aberrant neuronal activity has been reported in TDP-43 delta NLS mouse model of ALS, in which intrinsic hyperexcitability in layer V excitatory neurons of the motor cortex was observed. In contrast, this phenomenon was not present when the wild-type TDP-43 was expressed (15).

Topoisomerases are enzymes that regulate DNA supercoiling and are essential for proper neuronal activation (16). Topoisomerase 2 β (Top2β) is essential in mature neurons for the regulation of gene expression (16) and regulates activity-dependent gene expression in the brain through the generation of DSBs (12). Topoisomerases cleave positively supercoiled DNA ahead of transcription bubbles or replication forks (17). However, dysfunction of Top2β which stabilize the transient Top2β-DNA adduct are a substantial source of DNA damage (18). While etoposide, a Top2β poison (19), is widely used in studies of DNA damage, the role of TDP-43 in regulating Top2β function remains under investigated. Similarly, the role of TDP-43 protein in activity-dependent generation of DSBs remains unexplored.

In this study, we have determined for the first time the exact genomic locations of DSBs in neurons expressing ALS/FTD-associated TDP-43 mutants, A90V and A315T, compared to wild-type TDP-43 neurons. Interestingly, we found that DSBs occur within genes crucial for dysfunctional processes in ALS/FTD, including aberrant excitatory neurotransmission. Among these transiently induced breaks, we also identified persistent genomic targets, which were also detected *in vivo* in the TDP-43 rNLS mouse model. Furthermore, we identified two Top2β-dependent mechanisms that are disrupted by ALS/FTD-associated TDP-43 pathology, leading to the accumulation of DSBs. Specifically, we found that TDP-43 is required for the binding of Top2β to DNA, affecting both the stabilization of trapped Top2β-DNA cleavage complexes and/or the recruitment of Top2β to chromatin. Loss of TDP-43 function causes accumulation of supercoiled DNA, likely enhancing torsional stress and increasing susceptibility to DSB formation. Together, these findings uncover a previously unrecognized role of TDP-43 in regulating genome integrity during neuronal activation and implicate its dysfunction in activity-dependent DNA damage contributing to ALS/FTD pathogenesis.

## Results

### TDP-43 ALS/FTD-associated mutants render the neuronal genome more susceptible to DSBs than wild-type TDP-43

We employed the Breaks Labelling In Situ and Sequencing (BLISS) technique for the first time to investigate how two ALS/FTD-associated TDP-43 mutants, A90V and A315T, influence the formation of double-strand DNA breaks (DSBs) in mouse primary neurons. To achieve expression of these TDP-43 protein variants, neurons were transduced with lentiviruses encoding the respective TDP-43 constructs. Non-transduced neurons were eliminated through puromycin selection, as the lentiviral vector also included a puromycin resistance gene. Following selection, the remaining neurons underwent BLISS library preparation and sequencing. Analysis of DSB-specific genomic sites revealed that neurons expressing TDP-43 A90V mutant showed 6,876 unique gene targets, those expressing A315T exhibited 1054, while wild-type TDP-43 neurons had 1,483. Among the DSB-affected genes, we identified 766 genes commonly targeted in both A90V- and wild-type TDP-43 expressing neurons, and 183 genes shared between A315T and wild-type TDP-43 expressing neurons.

Given these overlaps, we next examined whether the number of unique DSBs per gene differs between mutants A90V or A315T and wild-type TDP-43. For this quantitative comparison we determined fold change of the unique DSBs loci in A90V or A315T samples relative to wild-type TDP-43 samples based on the normalized number of unique DNA breaks per gene. This comparison revealed that, although the same genes were targeted, neurons expressing A315T mutant exhibit a higher number of DSBs in most of the overlapping genes (131 genes, 71.6 % of total shared genes) compared to wild-type TDP-43 expressing neurons. Interestingly, neurons expressing A90V mutant were characterized by more DSBs in only 29 genes (3.8 % of total shared) compared to wild-type TDP-43 expressing neurons (**Figure 1** shows top 50 genes; full lists of genes in **Supplementary material 1**). The top 10 genes with the most pronounced increase in DSB frequency in A90V versus wild-type TDP-43 neurons were: *Csmd1, Plcb1, Arhgap26, Magi2, Rbms3, Ptprd, Igk, Cdh13, Opcml, Rora,* and in A315T vs wild-type TDP-43: *Zfhx3, Grip1, Kalrn, Ctdp1, Mpp2, Lmx1b, Cntn1, Fras1, Epn2, Ccdc33.* We additionally identified nine genes *Cdh13, Opcml, Spag16, Adcy2, Ncam2, Pard3b, Pcdh15, Gpc6, Alk* that were commonly targeted by DSBs in both A90V- and A315T-expressing neurons, but with less frequency in neurons expressing wild-type TDP-43.

**Figure 1.**
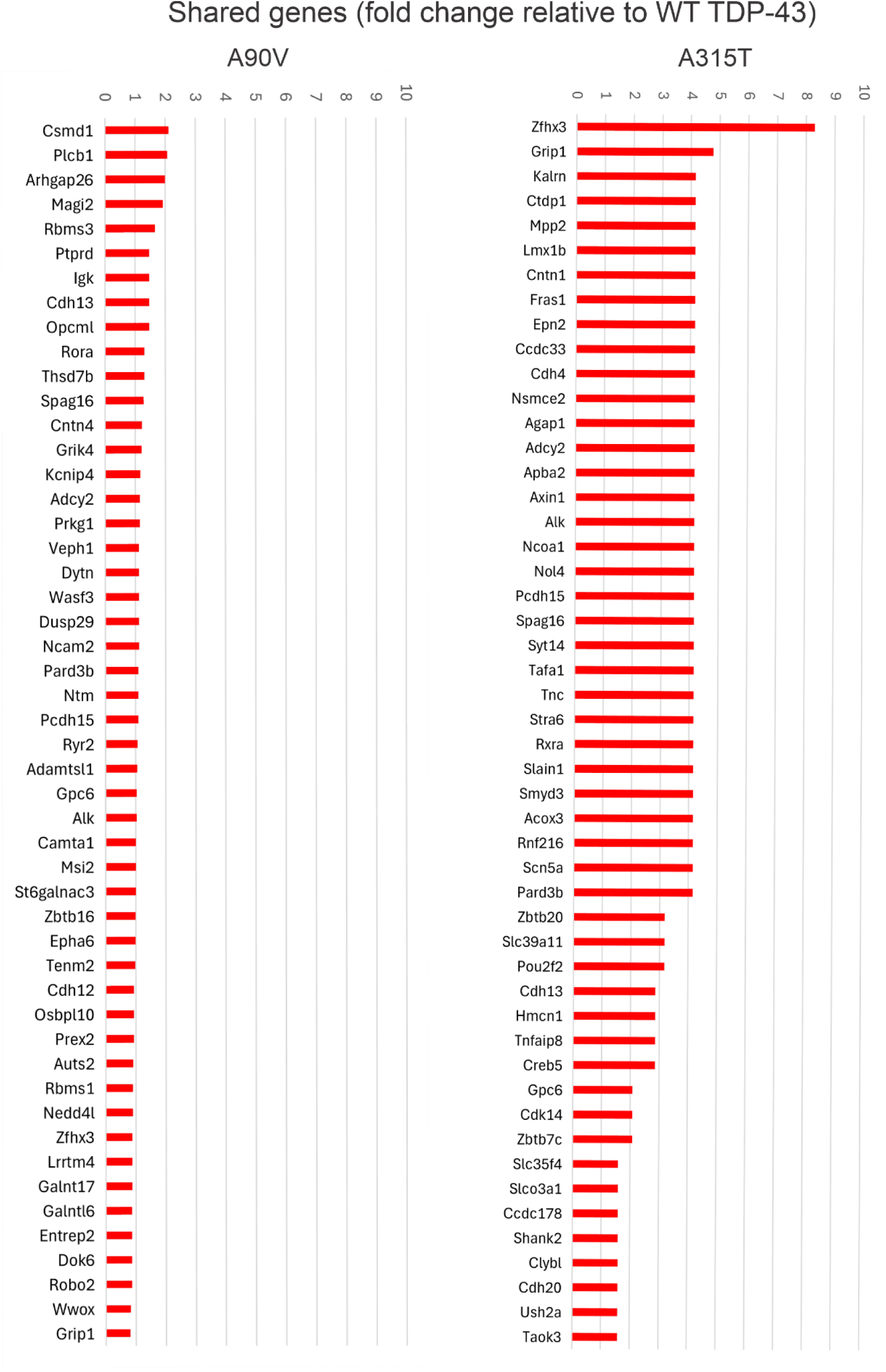
**Neurons expressing A315T or A90V TDP-43 mutants exhibit higher levels of DSBs within gene targets that are also shared with neurons expressing wild-type TDP-43**. The bars represent the fold change in the number of unique genomic targets of DSBs for each gene in for A90V (left) and A315T (right) samples relative to wild-type TDP-43 samples, shown for the top 50 gene targets. The data represent combined sequencing results from two independent runs of BLISS DNA libraries, performed on pooled TDP-43 variant samples prepared in three biological replications for each variant. Total number of pregnant mice used for the culture’s preparations = 6.

We then investigated genes uniquely affected by DSBs in neurons expressing either A90V or A315T mutants that were not targeted in wild-type TDP-43 neurons. We have identified 869 and 6110 such genes for A315T and A90V samples, respectively. The top ten uniquely targeted genes in A315T-expressing neurons were: *Unc5c, Mcph1, Kif2a, Ralgps1, Inpp4b, Pcdha4, Tprkb, Ugp2, Vgll4*, and *Vwa3a* and in A90V-expressing neurons: *Gpr158, Frmpd4, Lrp1b, Bcas3, Diaph3, Ros1, Dach1, Grid1, Dmd* and *Kazn* (full lists of genes **in Supplementary material 2**). Interestingly, we also identified 84 common genomic targets for neurons expressing A90V and A315T mutant (**Supplementary material 3**). The findings indicate that A315T and A90V mutants have a distinct impact on the magnitude and the range of the DSBs genomic targets. While A315T causes more severe damage to the shared DSB gene targets with wild-type TDP-43, A90V leads to DSBs in a larger number of genes compared to wild-type TDP-43.

### Excitatory neurotransmission is a significant target of DSBs in neurons expressing A90V TDP-43 mutant, but not wild-type TDP-43

Because neurons expressing the TDP-43 A315T mutant exhibited particularly high number of DSBs within the shared gene targets with wild-type TDP-43 neurons, and the A90V mutant were characterized with prominent increase of unique gene targets compared to both A315T and wild-type TDP-43 neurons, we focused our subsequent analysis on these sets of data. Thus, we performed a functional analysis using the ClueGO application to identify molecular functions associated with genes that were targeted in both A315T- and wild-type TDP-43- expressing neurons but were more heavily affected by DSBs in A315T-expressing cells. This approach revealed 7 statistically significant molecular functions, which were grouped into 4 functional categories. The most enriched category ‘structural constituent of postsynaptic density’ (42.86%), follow by ‘basal transcription machinery binding’ (28.57%), ‘cadherin binding’ (14.29%), and ‘signalling receptor complex adaptor activity’ (14.29%) (**Figure 2**).

**Figure 2.**
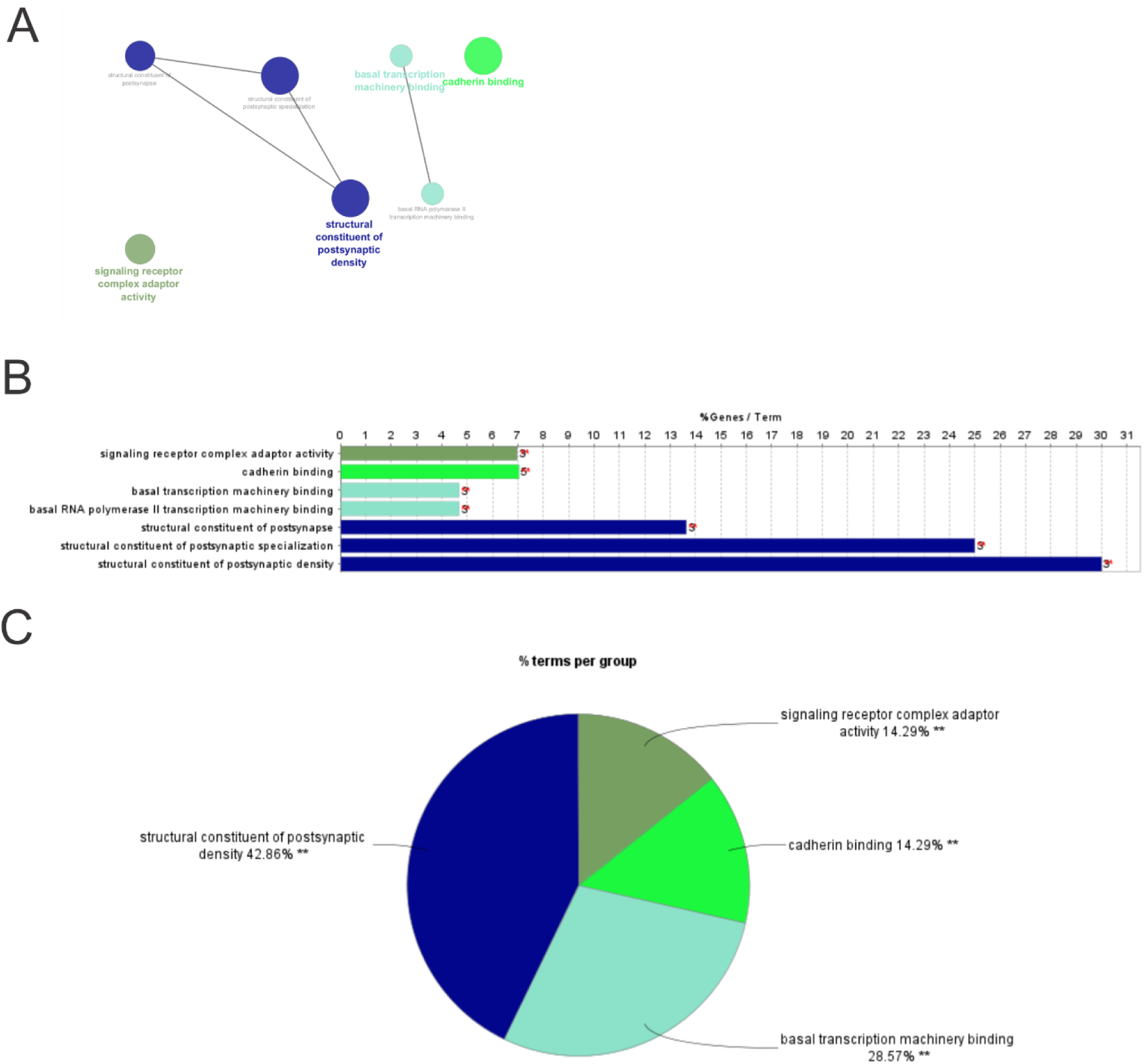
**Structural constituent of postsynaptic density is the most targeted functional category by DSBs in neurons expressing A315T TDP-43 mutant within shared DSBs gene targets with neurons expressing wild-type TDP-43 protein**. Functional analysis using the ClueGO application was performed to identify overrepresented molecular processes among the DSB gene targets common to both A315T mutant and wild-type TDP-43 neurons but targeted more often in A315T neurons. A The network representation of the 4 functional categories. The size of each node reflects the number of genes contributing to the corresponding molecular process. B The table shows 7 statistically significant molecular processes that are heavily targeted by DSBs in neurons expressing the A315T TDP-43 mutant compared to wild-type TDP-43 neurons. Group P Value < 0.01 corrected with Bonferroni step down. C The pie chart represents the percentage of terms in each functional category for 4 identified categories.

Next, we analysed the functional impact of DSBs on genes uniquely targeted in A90V-expressing neurons but not affected in neurons expressing wild-type TDP-43. This analysis identified 72 statistically significant molecular functions, grouped into 16 functional categories. The largest portion of these functions were ‘anion binding’ (29.63%) and ‘ion transmembrane transporter activity’ (24.69%). Notably, ‘glutamate receptor activity’, ‘ionotropic glutamate receptor activity’, and ‘glutamate receptor binding’ represented the most abundant molecular functions (**Figure 3 A and B**). This implies the excitatory neuronal transmission as a prominent target of DSBs specifically in neurons expressing the TDP-43 A90V mutant.

**Figure 3.**
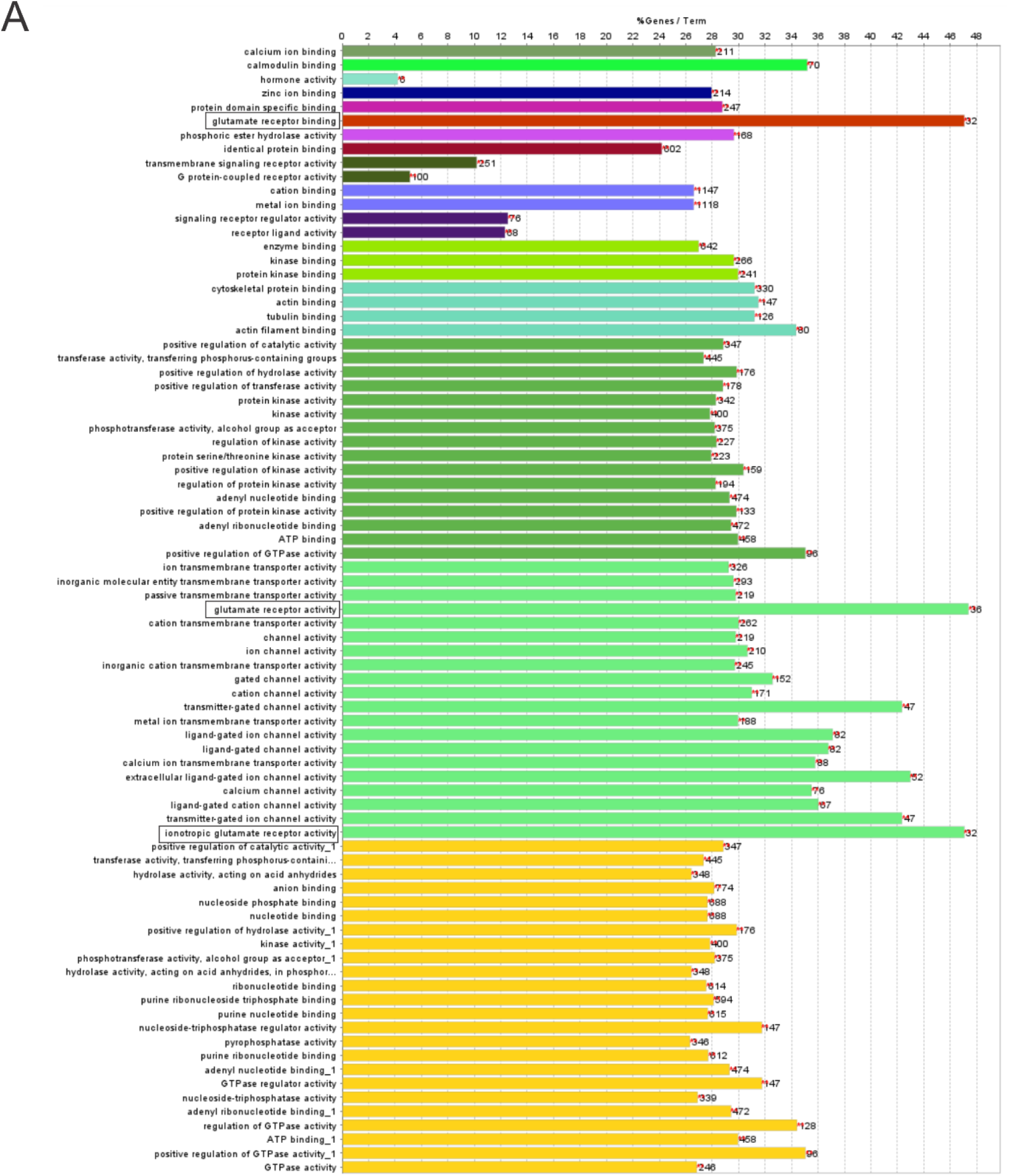
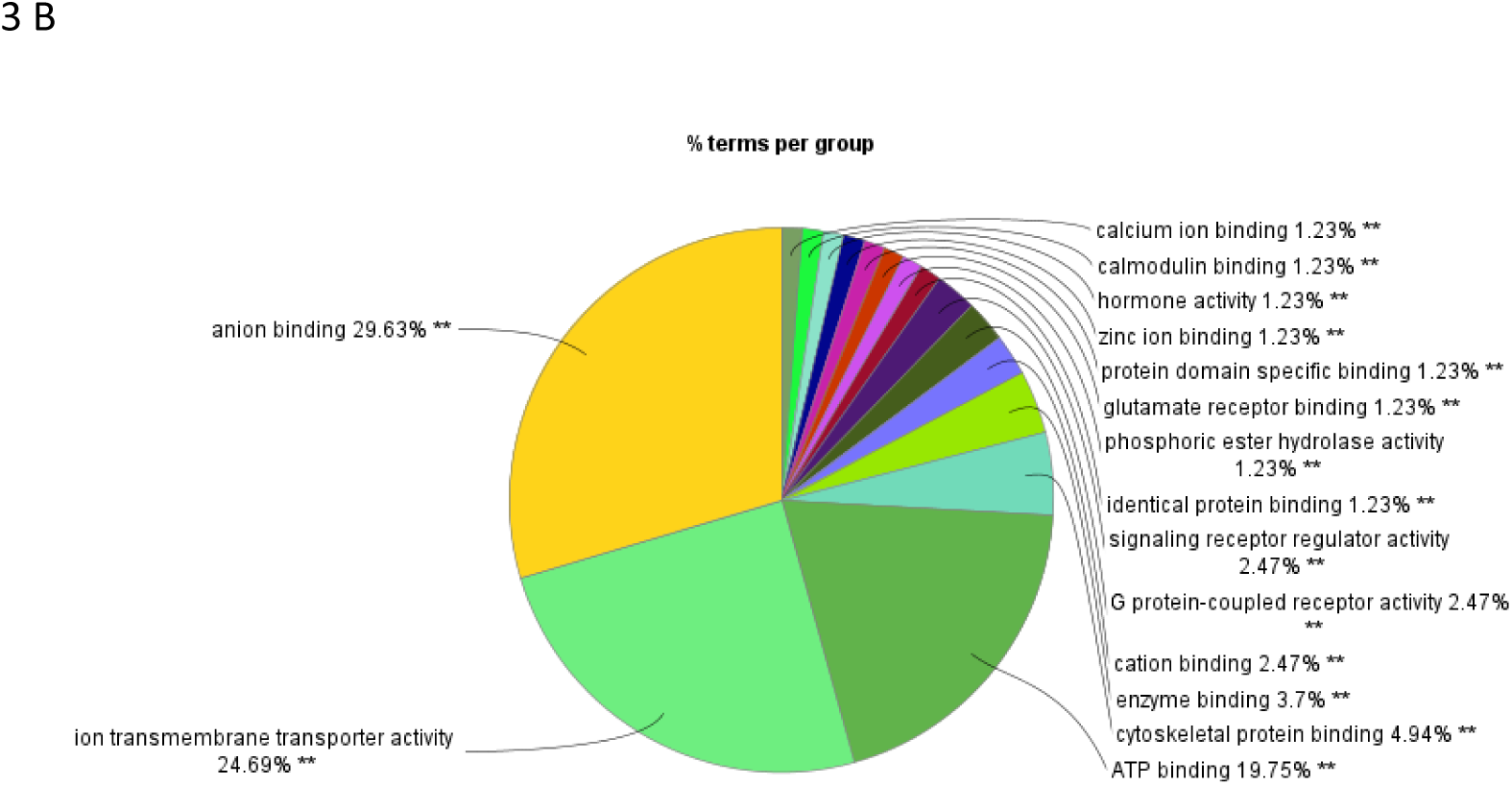
Excitatory neurotransmission is a prominent target of DSBs in neurons expressing the A90V TDP-43 mutant, but not in neurons expressing wild-type TDP-43. Functional analysis using the ClueGO application was performed to identify overrepresented molecular processes among the DSB gene targets present uniquely in neurons expressing A90V TDP-43 mutant. A The table shows 72 statistically significant molecular processes targeted by DSBs in neurons expressing A90V TDP-43 mutant. Group P Value < 0.001 corrected with Bonferroni step down. The ‘glutamate’ terms outlined in black frames. B The pie chart represents the percentage of terms in each functional category for 14 identified categories.

### DSBs caused by abnormal TDP-43 are both persistent and transient in nature

Given the possibility that some of the detected DSB loci may be transient due to ongoing DNA repair processes in neurons or target different regions each time, we investigated whether stable, recurrent genomic regions susceptible to DSBs exist in neurons expressing TDP-43 mutants. To address this, we employed qPCR, where the presence of DSBs at specific genomic sites is inferred from a reduction in amplification compared to undamaged DNA. We selected 10 genes for the analysis based on their links to ALS/FTD pathogenesis, roles in excitatory neurotransmission, and the extent of DSBs observed in our screen. We designed primers to amplify 400-1000 bp DNA fragments containing the DSBs identified by BLISS. We based on the assumption that DSBs may not always occur at the same genomic coordinates but are likely generated in nearby areas. Of these, we found the regions within 6 genes *Grm7, Gabbr2, Magi2, Cdk8, Sema3a, Gsk3b* that showed significantly reduced qPCR amplification in A90V-expressing neurons but not in those expressing wild-type TDP-43, indicating the presence of persistent DSBs at these loci. Next, we examined whether these regions are also targeted by DSBs *in vivo* using the TDP-43 rNLS mouse model of ALS (19). In this model, a mutation in the TDP-43 nuclear localization signal causes mis-localization of TDP-43 from the nucleus to the cytoplasm, leading to progressive neurodegeneration. Notably, one DSB target within the *Grm7* gene, which encodes the metabotropic glutamate receptor 7 (mGlu7), also showed a significant reduction in qPCR product in rNLS mice compared to age-matched controls (**Figure 4**). Interestingly, *Grm7* belongs to the set of genes targeted by DSBs in both A90V and A315T neurons, but not in wild-type TDP-43 neurons. Another target within the *Magi2* gene exhibited a clear trend toward reduced qPCR signal, although this did not reach statistical significance. Importantly, these analyses were performed approximately 2.5 weeks after doxycycline (Dox) withdrawal from mouse diet, a time point corresponding to early-stage pathology in this model (19). It is possible that DSB targets may emerge at more advanced disease stages. These findings suggest that a subset of stable DSB-prone genomic regions exists in neurons expressing the ALS/FTD-associated A90V TDP-43 mutant. However, the accumulation of a high number of DSBs in A90V neurons also points to a transient component, potentially contributing to functional impairment in these neurons.

**Figure 4.**
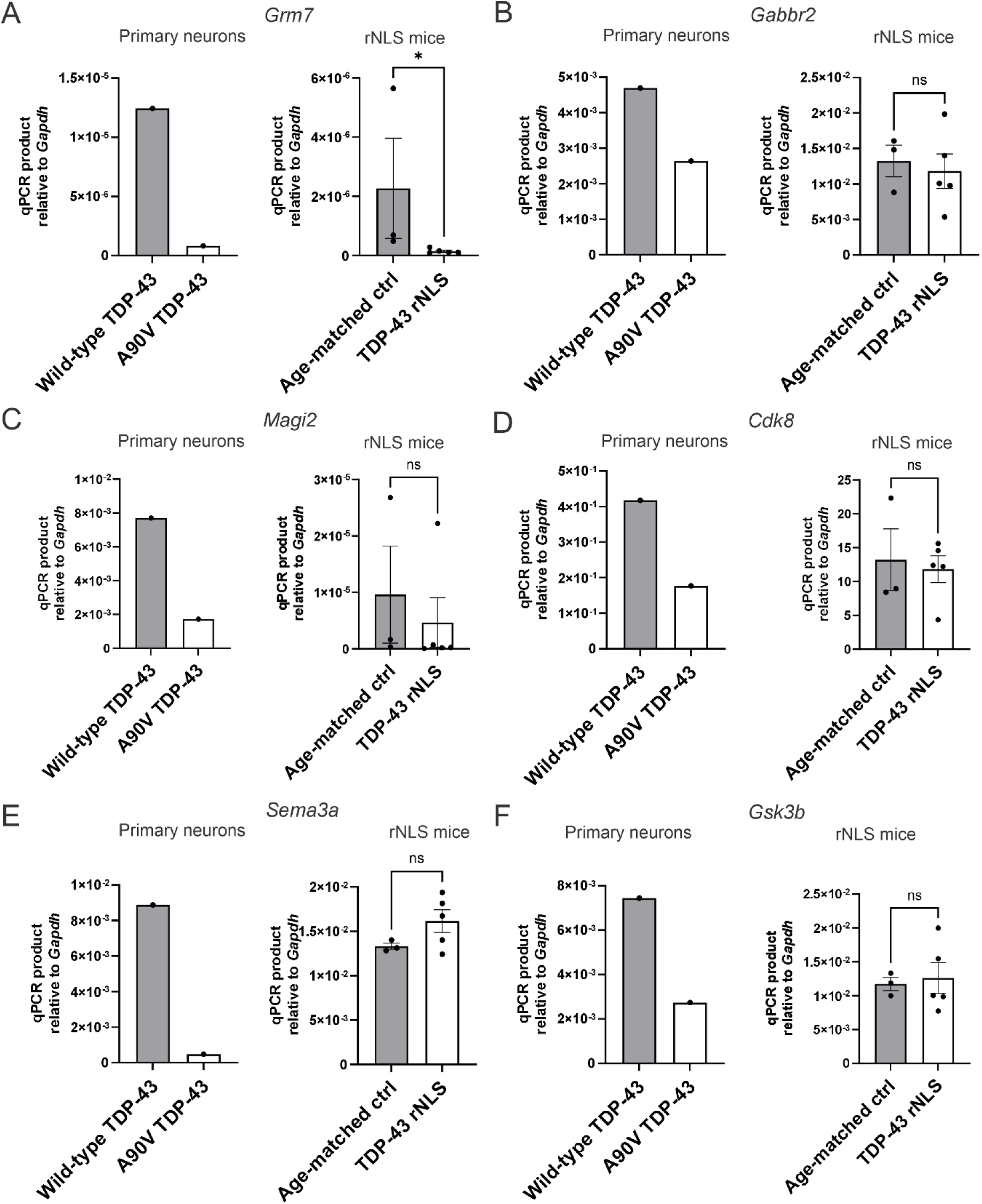
Selected genes with the confirmed regions targeted by DSBs in neurons expressing A90V TDP-43 mutant vs. wild-type TDP-43. qPCR was performed to target selected genomic coordinates identified by BLISS. A Grm7 is characterized with prominent reduction of amplicon in neurons expressing A90V TDP-43 mutant compared to wild-type TDP-43, indicating the presence of DNA breaks as well as in TDP-43 rNLS model compared to age-matched controls (t-test; *p < 0.05, 3 controls and 5 rNLS mice were used). B The reduction of Gabbr2 amplicon in neurons expressing A90V TDP-43 mutant compared to wild-type TDP-43 shows the presence of DNA breaks but not in TDP-43 rNLS model compared to age-matched controls (t-test; p < 0.05, 3 controls and 5 rNLS mice were used). C The redaction of Magi2 amplicon in neurons expressing A90V TDP-43 mutant compared to wild-type TDP-43 shows the presence of DNA breaks but not in TDP-43 rNLS model compared to age-matched controls. However, the trend towards reduction of the amplicon was observed in rNLS mice compared to age-matched controls. (t-test; p < 0.05, 3 controls and 5 rNLS mice were used). D The reduction of Cdk8 amplicon in neurons expressing A90V TDP-43 mutant compared to wild-type TDP-43 shows the presence of DNA breaks but not in TDP-43 rNLS model compared to age-matched controls (t-test; p < 0.05, 3 controls and 5 rNLS mice were used) E The reduction of Sema3a amplicon in neurons expressing A90V TDP-43 mutant compared to wild-type TDP-43 shows the presence of DNA breaks but not in TDP-43 rNLS model compared to age-matched controls (t-test; p < 0.05, 3 controls and 5 rNLS mice were used) F The reduction of Gsk3b amplicon in neurons expressing A90V TDP-43 mutant compared to wild-type TDP-43, shows the presence of DNA breaks but not in TDP-43 rNLS model compared to age-matched controls (t-test; p < 0.05, 3 controls and 5 rNLS mice were used).

### TDP-43 A90V mutant alters neuronal activation in topoisomerase 2β dependent manner

Given the excitatory neurotransmission is a target of DSBs, we next investigated whether A90V mutant alters neuronal activation, determining the expression of neuronal activation markers, c-Fos. Notable, c-Fos is an immediate early gene whose expression is regulated, in part, by DSBs generated by Topoisomerase IIβ (12). Under basal conditions, neurons expressing either wild-type or A90V TDP-43 exhibited robust c-Fos expression, with no significant differences observed between the two TDP-43 variants, as determined by both immunostaining and immunoblotting (**Figures 5**; immunostaining, p > 0.05; t-test, **Supplementary materials 6**). However, treatment with etoposide markedly reduced c-Fos expression in neurons expressing the A90V mutant compared to those expressing wild-type TDP-43, (**Figure 5**; immunostaining, ***p < 0.0001; t-test). Given that etoposide traps Top2β in DNA cleavage complexes, these results suggest that, unlike the A90V variant, wild-type TDP-43 may help prevent persistent Top2β-DNA trapping. This further implies that TDP-43 plays a role in neuronal activation pathways that rely on Top2β function.

**Figure 5.**
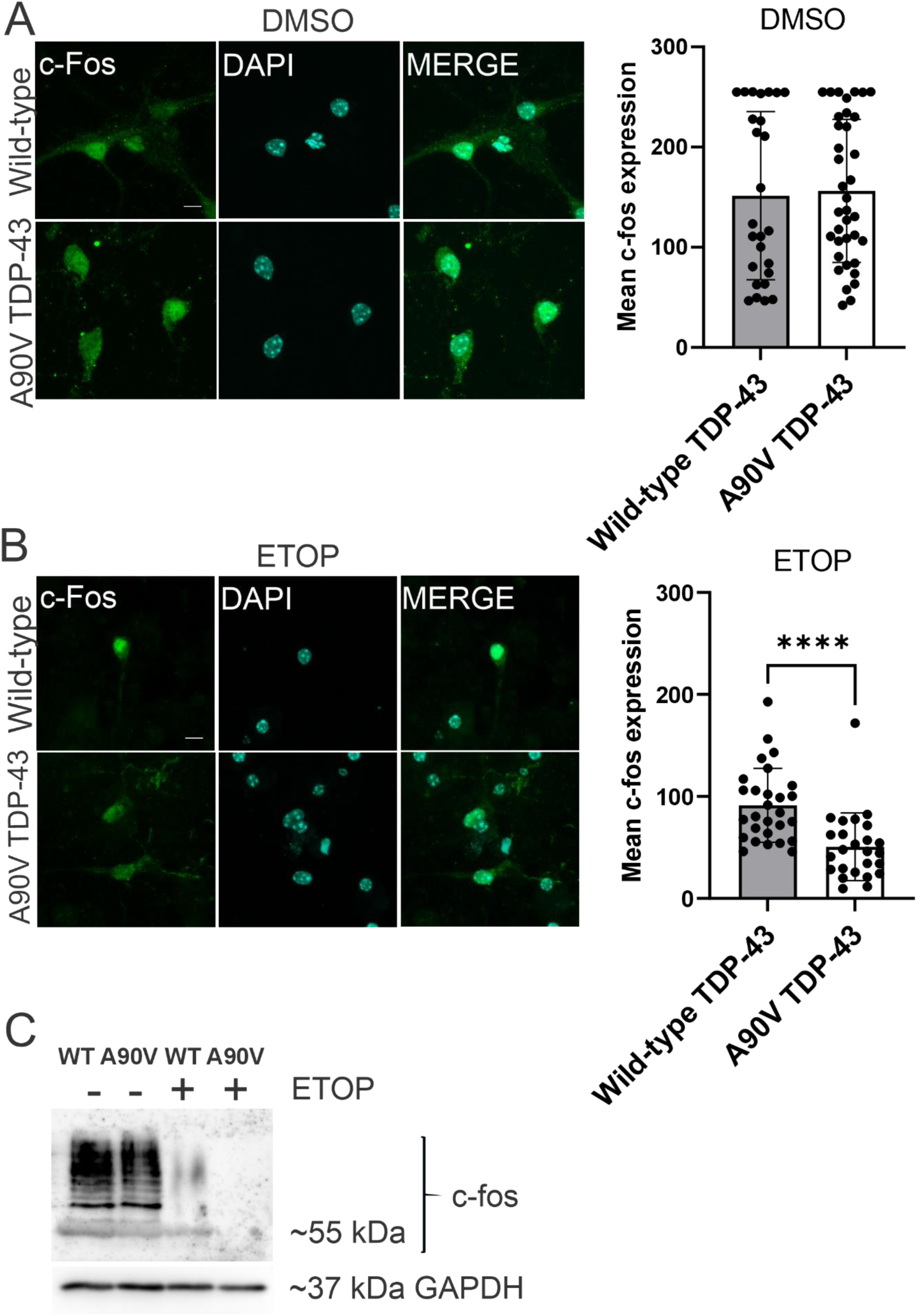
**Etoposide treatment impairs the expression of the neuronal activation marker c-Fos in neurons expressing the A90V TDP-43 mutant compared to wild-type TDP-43 neurons**. A Immunostaining for c-Fos in control, DMSO-treated mouse primary neurons expressing wild-type TDP-43 or the A90V TDP-43 mutant (representative images shown) and quantification of fluorescence intensity performed in Fiji revealed no significant differences in neuronal activation between these groups (t-test, p > 0.05, at least 24 neurons per group from three independent neuronal cultures were analysed; mean ± SD). B Immunostaining for c-Fos in etoposide-treated (ETOP) mouse primary neurons expressing wild-type TDP-43 or the A90V TDP-43 mutant (representative images shown), with quantification of fluorescence intensity performed in Fiji, revealed a statistically significant reduction of c-Fos expression in neurons expressing the A90V TDP-43 mutant compared to wild-type TDP-43 neurons. (t-test, ****p < 0.0001, at least 25 neurons per groups from three independent neuronal cultures were analysed; mean ± SD). C Immunoblotting for c-Fos was performed on lysates from neurons expressing wild-type TDP-43 or the A90V TDP-43 mutant following DMSO or etoposide (ETOP) treatment. ETOP treatment significantly impaired neuronal activation in A90V TDP-43-expressing neurons compared to wild-type TDP-43 neurons.

However, we anticipated differences in neuronal activation even in the absence of etoposide treatment as our BLISS data were obtained in such basal conditions. Since c-Fos is an immediate early gene with transient expression dynamics, we hypothesized that changes in neuronal activity might not be fully captured by assessing c-Fos expression alone. Therefore, we also examined the expression of phosphorylated CREB (pCREB), another marker of neuronal activity. Unlike c-Fos, pCREB acts as a long-lasting transcriptional activator, remaining active over extended periods and promoting the expression of genes that require prolonged transcriptional input (20). Determination of pCREB expression by immunostaining and immunoblotting revealed a statistically significant reduction in pCREB expression determined as immunofluorescence intensity in pCREB positive neurons expressing the A90V mutant compared to those expressing wild-type TDP-43 under basal conditions (DMSO-treated neurons; **Figure 6**; immunostaining, ****p < 0.0001; t-test). However, quantification of the percentage of neurons expressing pCREB and thus considered activated revealed that, although pCREB expression in neurons expressing the A90V TDP-43 mutant was reduced compared to wild-type TDP-43 neurons, a greater number of neurons became activated (**Figure 6**; * p < 0.005; t-test). This implies that wild-type TDP-43 neurons were capable for more effective activation likely in control synchronized way, while A90V TDP-43 mutant impairs this activation causing uncontrol asynchronized activation of larger fraction of neurons.

**Figure 6.**
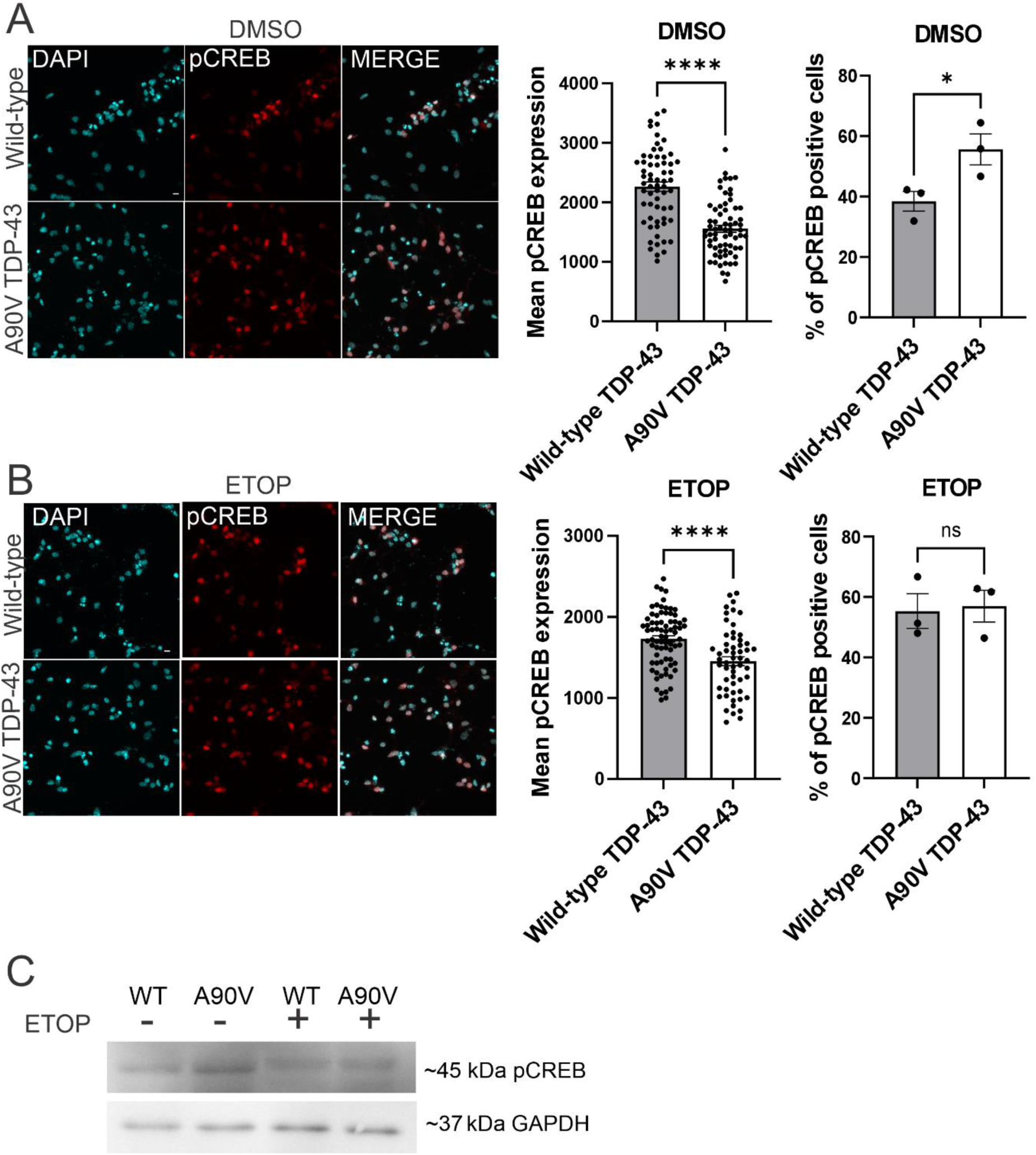
A90V TDP-43 mutant promotes prolong neuronal activation compared to wild-type TDP-**43.** A Representative images of immunostaining for pCREB in neurons expressing wild-type TDP-43 or the A90V TDP-43 mutant under control (DMSO-treated) conditions are shown. Quantification of signal intensity revealed a significant reduction of pCREB expression in neurons expressing the A90V TDP-43 mutant compared to wild-type TDP-43 (t-test, ****p < 0.0001; at least 63 neurons per group from three independent neuronal cultures; mean ± SEM). However, analysis of the percentage of pCREB-positive neurons showed an increase in the number of activated neurons expressing A90V TDP-43 relative to wild-type TDP-43 neurons (t-test, *p < 0.005; mean ± SEM). pCREB-positive neurons were quantified from three random fields per group using the Cell Counter plugin in Fiji. Scale bar 10 µm. B Representative images of immunostaining for pCREB in neurons expressing wild-type TDP-43 or the A90V TDP-43 mutant after etoposide treatment (ETOP-treated) are shown. Quantification of the immunofluorescent signal intensity revealed decrease in pCREB expression in neurons expressing A90V TDP-43 mutant compared to wild-type TDP-43 (t-test; **** p < 0.0001; at least 56 neurons per group from three independent neuronal cultures were analysed; mean ± SEM). Quantification of pCREB-positive neurons revealed no differences in the percentage of activated neurons between those expressing A90V and wild-type TDP-43 (t-test, p > 0.05; mean ± SEM). pCREB-positive neurons were quantified from three random fields per group using the Cell Counter plugin in Fiji. Scale bar 10 µm. C Immunoblotting for pCREB performed on neurons expressing A90V or wild-type TDP-43 and treated with DMSO or etoposide (ETOP) showed a visible increase in pCREB expression in A90V TDP-43-expressing neurons compared to wild-type TDP-43 under control (DMSO-treated) conditions.

Treatment with etoposide led to a significantly greater reduction of pCREB expression determined as immunofluorescence intensity in neurons expressing the TDP-43 A90V mutant compared to those expressing wild-type TDP-43 (**Figure 6**; immunostaining, ****p < 0.0001, t-test). However, there was no statistically significant difference in the percentage of pCREB-positive neurons between the groups investigated (**Figure 6**; immunostaining; p > 0.05, t-test). Furthermore, immunoblotting for pCREB revealed an increase in pCREB levels in neurons expressing the A90V TDP-43 mutant compared to those expressing wild-type TDP-43 under basal conditions. No noticeable differences in pCREB expression between A90V and wild-type TDP-43 neurons were observed following etoposide treatment. These findings suggest that the immunoblotting results likely reflect a higher number of pCREB-positive cells in the A90V TDP-43 condition, rather than an increase in the intensity of pCREB expression per cell (**Figure 6 C**, **Supplementary materials 6**).

The findings imply that TDP-43 ALS/FTD associated mutant, A90V alters neuronal ability for activation and potentially impact both expression of early response genes and genes required prolong transcriptional activation.

### Overexpression of tyrosyl-DNA phosphodiesterase 2 (TDP2) alters neuronal activation and prevents DNA damage in A90V mutant neurons compared to A90V mutant without TDP2 overexpression

To gain deeper insight into the observed phenomenon, we next investigated whether tyrosyl-DNA phosphodiesterase 2 (TDP2), an enzyme that removes covalently trapped Top2β at DNA is involved in the mechanism of altered neuronal activation in neurons expressing A90V TDP-43 mutant. To test this, neurons expressing the ALS-associated TDP-43 A90V mutant were additionally transduced with lentiviral vectors encoding a doxycycline-inducible *Tdp2*. Then, we assessed c-Fos expression by immunoblotting in neurons with and without Tdp2 overexpression (doxycycline treated vs. untreated). Interestingly, Tdp2 overexpression enhances c-Fos expression in A90V-expressing neurons compared to non-doxycycline-treated controls, indicating the involvement of Tdp2 in neuronal activation. In contrast, Tdp2 overexpression led to a reduction in pCREB levels compared to non-doxycycline-treated controls (**Figure 7, Supplementary materials 6**), implying a differential regulatory effect on activation-related signalling pathways.

**Figure 7.**
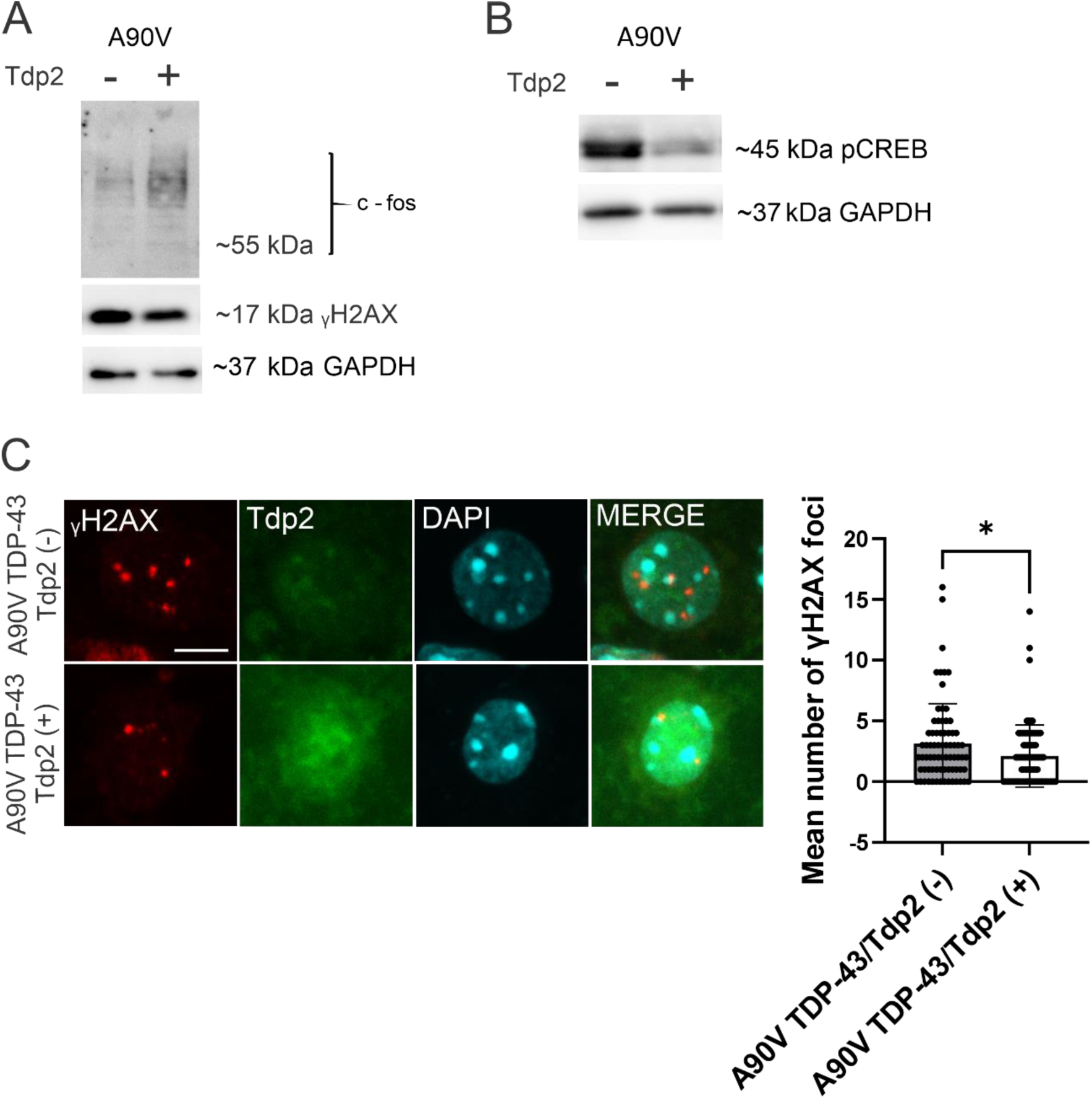
**Tyrosyl-DNA phosphodiesterase 2 (TDP2) reduces DNA damage and facilitates neuronal activity in A90V TDP-43 mutant neurons**. A Immunoblotting revealed that overexpression of TDP2 enhanced c-Fos expression and reduced γH2AX levels in neurons expressing the A90V TDP-43 mutant compared to neurons expressing the A90V TDP-43 mutant without TDP2 overexpression. B The expression of TDP2 was induced by doxycycline treatment. In contrast to c-Fos, pCREB expression was reduced by TDP2 overexpression in neurons expressing the A90V TDP-43 mutant compared to control (non-overexpressing) A90V TDP-43 neurons, determined by immunoblotting. C Quantification of γH2AX foci confirmed a reduction in DNA damage in neurons expressing the A90V TDP-43 mutant following TDP2 overexpression compared to control A90V TDP-43 neurons (t-test, *p < 0.05, at least 75 neurons per group from three independent neuronal cultures were analysed; mean ± SD). Scale bar 10 µm.

Then, we assessed DNA damage using the DNA damage marker, γH2AX. Tdp2 overexpression visibly reduced DNA damage levels in A90V-expressing neurons compared to those without Tdp2 induction, detected with immunoblotting and immunostaining (**Figure 7**; *p < 0.05; t-test). These findings imply that altered neuronal activation is coupled with DNA damage in neurons expressing the A90V mutant.

However, since Tdp2 overexpression only partially alleviates DSB accumulation, we expected that, in addition to trapped Top2β, other mechanisms also contribute to DSB formation in the context of TDP-43 A90V pathology.

### TDP-43 knock down impairs Top2β binding to DNA leading to the accumulation of supercoiled DNA

To determine whether TDP-43 protein can impact Top2β function in another way than trapping Top2β pathological complexes and thus leading to the extensive DSBs, we first performed a modified Rapid Approach to DNA Adduct Recovery (RADAR) assay on mouse primary cortical neurons after TDP-43 knockdown using specific siRNA versus an off-target control. Initially, we validated the efficiency of siRNA-mediated knockdown by immunostaining and immunoblotting. Both immunostaining and immunoblotting confirmed effective TDP-43 knockdown in mouse primary neurons (**Figure 8 A and B, Supplementary materials 6**).

**Figure 8.**
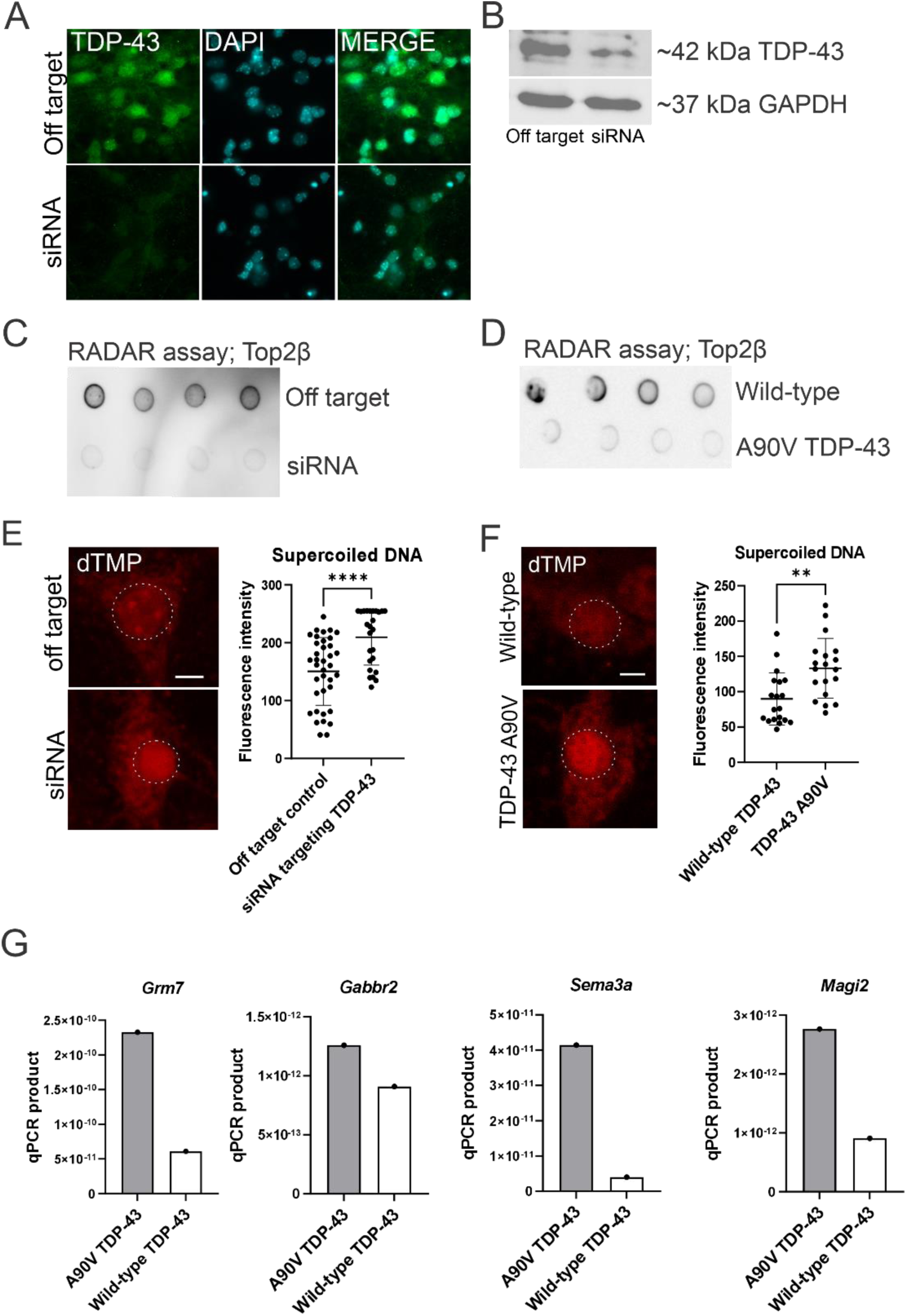
**TDP-43 depletion or A90V mutant expression impairs Top2β binding and leads to the accumulation of supercoiled DNA in mouse primary neurons**. **A** Validation of TDP-43 knockdown efficiency in mouse primary neurons was performed by immunostaining for TDP-43. Compared to the off-target control, TDP-43 knockdown using specific siRNA effectively reduced TDP-43 expression. **B** Validation of TDP-43 knockdown efficiency in mouse primary neurons was performed by immunoblotting for TDP-43, which confirmed a reduction of TDP-43 expression in neurons treated with specific siRNA compared to the off-target control. **C** The representative RADAR assay shows a marked reduction of Top2β bound to DNA in neurons with TDP-43 knockdown using specific siRNA compared to the off-target control. Four dots in each row represent technical replicates, with a total of three biological replicates performed. **D** The representative RADAR assay shows a marked reduction of Top2β bound to DNA in neurons expressing TDP-43 A90V mutant compared to wild-type TDP-43. Four dots in each row represent technical replicates, with a total of three biological replicates performed. **E** Intercalation of psoralen conjugated to biotin (dTMP) into supercoiled DNA revealed a significantly higher amount of supercoiled DNA in neurons with TDP-43 knockdown using specific siRNA compared to off-target control neurons. Left panel, representative images showing dTMP signal detected with streptavidin-DyLight 550 labelling. Right panel, the signal intensity quantification using Fiji (ImageJ) (****p < 0.0001, t-test, at least 25 neurons per groups from three independent neuronal cultures were analysed; mean ± SD). Scale bar 10 µm. **F** Intercalation of psoralen conjugated to biotin (dTMP) into supercoiled DNA revealed a significantly higher amount of supercoiled DNA in neurons expressing TDP-43 A90V mutant compared to wild-type TDP-43. Left panel, representative images showing dTMP signal detected with streptavidin-DyLight 550 labelling. Right panel, the signal intensity quantification using Fiji (**p < 0.001, t-test; at least 19 neurons per group from three independent cultures were analysed; mean ± SD). Scale bar 10 µm. **G** qPCR analysis revealed that the formation of supercoiled DNA containing DSB gene targets (Grm7, Gabbr2, Sema3a, and Magi2) is enhanced in A90V TDP-43 neurons compared to wild-type TDP-43 neurons.

For the RADAR assay, DNA extracted from TDP-43 knockdown and control neurons was spotted at equal concentrations onto a nitrocellulose membrane, followed by immunoblotting with an anti-Top2β antibody. The intensity of the Top2β signal reflects the amount of Top2β bound to DNA. TDP-43 knockdown with specific siRNA markedly reduced the binding of Top2β to DNA compared to the off-target control in mouse primary neurons, indicating that TDP-43 is required for efficient Top2β and DNA binding (**Figure 8 C, Supplementary materials 6**). Given that Top2β prevents torsional stress arising due to the helical structure of DNA, we propose that TDP-43 knockdown may lead to the accumulation of unresolved supercoiled DNA. Consequently, this may increase torsional stress and place DNA at greater risk of DSBs. To test this possibility, we assessed the accumulation of supercoiled DNA in neurons with TDP-43 knockdown compared to off-target controls. Therefore, we used psoralen conjugated with biotin (dTMP), which preferentially intercalates into supercoiled DNA (21). Quantification of the fluorescence signal using Fiji software revealed a statistically significant increase in the accumulation of supercoiled DNA in neurons with TDP-43 knockdown compared to off-target control neurons (****p < 0.0001, *t*-test) (**Figure 8 E**). These findings imply that TDP-43 is required for Top2β binding to DNA, thereby enabling Top2β to resolve supercoiled DNA and prevent torsional stress, a potential cause of DSB formation.

### TDP-43 A90V mutant, but not wild-type TDP-43 impairs Top2β binding to DNA leading to the accumulation of supercoiled DNA

Next, we investigated whether the TDP-43 A90V mutant impairs Top2β binding to DNA and thereby contributes to the accumulation of supercoiled DNA. To this end, we performed the modified RADAR assay in mouse primary neurons expressing either the TDP-43 A90V mutant or wild-type TDP-43. Similar to the effect observed with siRNA-mediated TDP-43 knockdown, the A90V mutant significantly reduced Top2β binding to DNA compared to wild-type TDP-43, indicating that TDP-43 mutant disrupts the interaction between Top2β and DNA (**Figure 8 D, Supplementary materials 6**). We then quantified the levels of supercoiled DNA using biotin-conjugated dTMP, as described above. Neurons expressing the TDP-43 A90V mutant showed a significantly higher accumulation of supercoiled DNA than those expressing wild-type TDP-43 (**p < 0.001, t-test) (**Figure 8 F**). These findings demonstrate that the TDP-43 A90V mutant, unlike the wild-type TDP-43 protein, interferes with Top2β-DNA binding and leads to the accumulation of DNA supercoiling.

To further investigate whether the accumulation of supercoiled DNA and the resulting torsional stress contribute to extensive DSB formation in neurons expressing the TDP-43 A90V mutant, we examined whether regions of supercoiled DNA overlap with DSB genomic targets identified by BLISS. To assess this, we used biotin-conjugated dTMP, which intercalates into supercoiled DNA in neurons expressing the A90V mutant and wild-type TDP-43. Supercoiled DNA fragments were then selectively isolated using streptavidin-coated beads, followed by qPCR amplification of regions corresponding to DSB genomic targets identified by BLISS. Thus, we designed primers targeting nearby location of DSB sites identified previously within genes *Grm7*, *Magi2, Sema3a, Gsk3b, Gabbr2* (Figure 5) and generating short 59 bp product. The *Cdk8* gene was excluded from the investigation, as it was not possible to design accurate primers due to the high content of GG/CC sequences. Consistent with our findings, we were able to pull down supercoiled DNA in A90V TDP-43 samples, in contrast to wild-type TDP-43, confirming that neurons expressing A90V TDP-43, but not wild-type TDP-43 are reach in supercoiled DNA. Although DNA was not detectable in the wild-type TDP-43 samples, we use them as a negative control in qPCR. Among the five investigated sequences, three *Grm7*, *Sema3a*, and *Magi2* were clearly more abundant in A90V TDP-43 samples, and one *Gabbr2* was slightly more abundant compared to wild-type TDP-43 samples, indicating enhanced formation of supercoiled DNA containing these genes in A90V samples relative to the wild-type TDP-43. The gene *Gsk3b* was not detectable in any of the samples (not shown), suggesting that causes other than supercoiling may be responsible for the DSBs observed within this gene (**Figure 8 G**). However, we cannot exclude the possibility that *Gsk3b* was not detectable due to limitation of the method. The findings support the notion that, in contrast to wild-type TDP-43, the A90V variant promotes the accumulation of supercoiled DNA in neurons, which may facilitate the generation of DSBs.

## Methods

### Transgenic mice

Transgenic bigenic rNLS mice and their monogenic littermate controls were generated by breeding two distinct monogenic lines: B6;C3-Tg(NEFH-tTA)8Vle/J (NEFH-tTA line 8, stock no. 025397) and *B6;C3-Tg(tetO-TARDBP)4Vle/J*\* (tetO-hTDP-43ΔNLS line 4, stock no. 014650). Both lines were maintained on a mixed B6/C3H genetic background. Breeding pairs were originally obtained from The Jackson Laboratory (USA), and genotyping was performed following the supplier’s standard protocols. To suppress expression of the hTDP-43 transgene, doxycycline was incorporated into the diet at a concentration of 200 mg/kg (Gordon Specialty Feeds).

### Primary neuronal culture

The timed pregnant mice were obtained from Flinders University Animal Facility. Primary cortical neurons were isolated from E16 C57BL/6 mouse brains and plated onto glass coverslips coated with poly-L-lysine (Sigma-Aldrich, #P2636-25MG). The neurons were cultured in media containing B27 (Gibco, cat # 17504044), neurobasal medium (Gibco, #10888022), 1% GlutaMAX (Gibco, # 35050061), and 1% penicillin/streptomycin (Gibco, #15140122). The cultures were maintained at 37°C with 5% CO^2^.

### Lentiviruses and transgenes expression in mouse primary neurons

The viruses have been ordered from Gene Silencing and Expression Core Facility, Adelaide University, South Australia. The following viral vectors were used pLVX-Puro-TDP-43-A315T, which was a gift from Shawn Ferguson (Addgene plasmid #133755; https://www.addgene.org/133755/; RRID: Addgene_133755). pLVX-Puro-TDP-43-WT, which was a gift from Shawn Ferguson (Addgene plasmid # 133753; https://www.addgene.org/133753/; RRID: Addgene_133753). pLVX-Puro-TDP-43-A90V, which was a gift from Shawn Ferguson (Addgene plasmid #133754; http://n2t.net/addgene:133754; RRID:Addgene_133754). The pLV[Exp]-EGFPTRE> hTDP2[NM_016614.3] (VB230713-1608tzp) was purchased from VectorBuilder. 24h after lentiviral infection to express TDP-43 variants, the mouse primary neurons were treated with 250ng of puromycin overnight to eliminate non-transduced neurons. To co-express TDP2, neurons were additionally infected with suitable viruses and treated with 50ng doxycycline overnight to induces TDP2 expression. Because TDP2 expression is indicated by EGFP fluorescence, successful expression of TDP2 was confirmed by microscopy.

### TDP-43 knock down with specific siRNA

ON-TARGET plus Mouse Tardbp (230908) siRNA - SMART pool (# L-040078-01-0005) and ON-TARGET plus non targeting pool (# D-001810-10-05) were purchased from Horizon Discovery. Neurons were transfected with siRNAs pool or control pool using Lipofectamine 2000 (Invitrogen). Experiments were performed 48 h after transfection.

### A modified ‘rapid approach to DNA adducts recovery’ (RADAR) assay

Mouse primary neurons were lysed with RLT buffer (Qiagen) followed by DNA precipitation with 3M sodium acetate and ethanol and centrifuged at 20000g. After centrifugation the pellet was resuspended in 8 mM NaOH with heating at 95°C. DNA concentration was determined with Nanodrop spectrophotometer. Then 200 ng of DNA in TE buffer per sample was spotted on nitrocellulose membrane followed by blocking with 5% Skim Milk for 1h and incubation with primary antibody rabbit anti-Top2β (1:1000; Invitrogen #PA5-27750) overnight at 4°C. After 3x washing with 1xTBS buffer the membrane was incubated with goat anti-rabbit IgG (H+L) secondary Antibody, HRP conjugated (1:1000, Invitrogen, #31460) and the membrane was imaged with Image Quant LAS 2000 (GE Healthcare). The expression of Top2β signal was measured with Fiji software (22).

### Immunoblotting

Neurons were lysed in RIPA buffer (Triton X-100, 1%; Sodium Deoxycholate, 1%; SDS, 0.1%; NaCl, 150 nM; Tris, pH 7.4, 10 mM; PMSF, 1mM) supplemented with protease inhibitor cocktail (cOmplete™ ULTRA Tablets, Mini, EASYpack, Roche 05892970001). A BCA protein assay (Pierce) was employed to quantify protein concentrations of each sample. SDS-PAGE was then performed, followed by blotting onto nitrocellulose membranes (Bio-Rad). Blocking was performed using 5 % skim milk then incubated with primary antibodies: rabbit anti-cFOS (K-25) (1:500, Santa Cruz, #E1606), rabbit anti-p-CREB (1:500, Cell Signalling, #9198S), mouse anti-GAPDH (1:2000, Invitrogen, # AM27828), rabbit anti-γH2AX (1:300, Betyl, #A700-053) overnight in blocking buffer at 4 °C. The blots were incubated with HRP-conjugated secondary antibodies (goat anti-rabbit or anti-mouse IgG (1:1000, Invitrogen, #31460 and #31430) for 1–2 h at room temperature following 3x washing with 1xTBS buffer. Images were obtained following development using ECL blotting substrates (Bio-Rad) with a Image Quant LAS 2000 (GE Healthcare).

### Immunocytochemistry

Mouse primary neurons were fixed in 4% paraformaldehyde for 10 min followed by 3 x washing with 1 x PBS. The blocking was performed in 5% BSA with 0.1 % Triton and incubated with primary antibodies at 4°C overnight. The following primary antibodies were used: rabbit anti-cFOS (K-25) (1:200, Santa Cruz, #E1606), rabbit anti-pCREB (1:200, Cell Signalling, #9198S), rabbit anti-γH2AX (1:200, Betyl, #A700-053) or mouse anti-TDP-43 (1:500, Proteintech, #66734-1-Ig). After 3x washing with 1xPBS the secondary antibody was applied. The following secondary antibodies were used: anti rabbit or anti mouse Alexa Fluor 488 or 568 (1:500, Invitrogen, #A11008, #A11001, #A11004, #A11011). The nuclei were counterstained with DAPI during second wash with 1xPBS.

### DNA damage induction

DNA damage in neurons was induced with etoposide at 13.5 uM concentration as described previously (6).

### BLISS DNA library preparation

BLISS DNA library was prepared according to the previously described protocol (23). Briefly, neurons were crosslinked on plates with 2% paraformaldehyde, followed by treatment with 125 mM glycine and a wash with ice-cold 1× PBS. The neurons were then subjected to lysis with the first lysis buffer (10 mM Tris-HCl, 10 mM NaCl, 1 mM EDTA, 0.2% Triton X-100, pH 8.0) at 4 °C for 60 minutes to remove the cytoplasm, followed by lysis with the second lysis buffer (10 mM Tris-HCl, 150 mM NaCl, 1 mM EDTA, 0.3% SDS, pH 8.0) at room temperature for nuclear permeabilization. The neurons were then subjected to *in situ* double-strand break (DSB) blunting using the Quick Blunting Kit (NEB #E1201), followed by overnight ligation of BLISS adapters using T4 ligase. The following day, unlighted adapters were washed off with a 1 M high-salt buffer and rinsed with water prior to DNA extraction. Nuclei were lysed in Tail buffer (10 mM Tris, 100 mM NaCl, 50 mM EDTA, 1% SDS, pH 7.5) supplemented with proteinase K and incubated at 55 °C overnight. DNA was extracted using phenol/chloroform/isoamyl alcohol, DNA concentrations were equalized and subjected to shearing with a Bioruptor Pico (Diagenode) to obtain 350–600 bp fragments. DNA obtained from three independent cultures of primary neurons, delivered from three mice and constituted 3 biological replications for each TDP-43 variant, was pooled. AMPure XP beads (Beckman Coulter, #A63880) were used to concentrate the DNA, and DNA concentration was measured using a Qubit fluorometer. An equal amount of DNA was used for BLISS library preparation, which included the following steps: *in vitro* transcription using the MEGAscript T7 Kit (Thermo Fisher, #AM1334), RNA purification with RNA Clean XP beads (Beckman Coulter, # A63987), Illumina RA3 adapter ligation with T4 ligase highly conc. 5U/ul (Thermo #EL0011), reverse transcription with SuperScript IV Reverse Transcriptase (Thermo Fisher Scientific, #18090200), and library indexing amplification by PCR using the NEBNext Ultra II Q5 Master Mix (New England Biolabs, #M0544L). In the final step, the library was size-selected and purified using AMPure XP beads (Beckman Coulter, #A63880). The samples were sent to the Australian Genome Research Facility (AGRF) for quality check and sequencing, which was performed on a NovaSeq X system using a 10B lane and 300-cycle run. Two independent replications of this experiment, each consisting of pooled samples and performed in separate sequencing runs, were conducted. The primers and adapters used in this protocol were ordered from Integrated DNA Technologies and are listed in **Supplementary materials 4**.

### Quantitative PCR (qPCR)

The assessment of the presence of DSBs were done based on previously described protocol (24). Genomic DNA from neurons was extracted using Tail buffer (10 mM Tris, 100 mM NaCl, 50 mM EDTA, 1% SDS, pH 7.5) and proteinase K digestion, followed by phenol/chloroform/isoamyl alcohol extraction. Mouse brains were snap-frozen in TRIzol reagent, and genomic DNA was subsequently isolated using the phenol/chloroform/isoamyl alcohol method. For qPCR analysis, 10 ng of genomic DNA was mixed with PowerUp™ SYBR™ Green Master Mix (Applied Biosystems, #A25742) and the appropriate primers (listed in **Supplementary materials 5**). Each sample was analysed in duplicate on Rotor-Gene Qiagen thermocycler under the following conditions, for DSB detection: an initial denaturation at 95°C for 3 min, followed by 35 cycles of 94 °C for 1 min, annealing at the primer-specific temperature for 1 min, and 72 °C for 1 min, final incubation of 4 min at 72°C. The housekeeping gene *Gapdh* was used as an internal control with the condition: 95°C for 3 min, followed by 35 cycles of 94°C for 1 min, 60°C for 1 min, and 72 °C for 1 min, hold at 72°C for 4 min. A non-template control was included in each run. Relative amplicon abundance was determined by normalizing to *Gapdh*, and fold changes were calculated using the 2^–ΔΔCt method (25). For the detection of the sequences within supercoiled DNA in A90V and wild-type TDP-43 samples following conditions were used: hold at 60°C for 1 min, then 40 x cycles: 95°C for 20 secs, 58°C for 20 secs, 72°C for 20 secs, hold at 60°C for 1 min. The same data analysis approach used for DSB assessment was applied; however, the amplicon abundance of A90V samples was compared to that of control wild-type TDP-43 samples.

### BLISS sequencing data post-processing

The BLISS sequencing data were analysed with the pipeline publicity available and described previously (23). Thus, the R1 reads obtained from sequencing were processed through the pipeline, allowing for 1 mismatch during UMI filtering process, yielding the genomic coordinates of DSBs. The obtained genomic coordinates were annotated to the mouse reference genome (GRCm39) to determined which genes are associated with DNA breaks using bedtool intersect tool. The custom python algorithm was used for the comparative analysis of overlapping DSBs targets between investigated groups (available at https://github.com/AnnaEKonopka/DSBs_geneID_comparison-.git). For quantitative comparison of DNA damage-strand breaks (DSBs) the number of unique molecular identifiers (UMIs) per gene was first normalized by the total number of DSB ends detected in each sample for each biological replication. This normalization accounts for differences in sequencing depth between samples. The normalized values for each replication were subsequently averaged per gene and use to calculate the fold changes between TDP-43 mutants and wild-type TDP-43 samples. The open-source ClueGO analysis was performed on the identified gene lists to determine overrepresented pathways.

### Supercoiled DNA investigation (Psoralen assay)

Neurons were treated with 100 µg/ml dTMP (EZ-Link™ Psoralen-PEG3-Biotin, Thermo Scientific, #29986) for 10 min at room temperature in the dark. Subsequently, bTMP was cross-linked to DNA by exposure to 360 nm UV light for 10 min at room temperature. The neurons were then subjected to either fluorescent labelling or supercoiled DNA precipitation assays. For fluorescent labelling, neurons were fixed with 4% paraformaldehyde (PFA), washed with three times with 1xPBS, and blocked with 5% BSA for 1 h at room temperature. Cells were then incubated with streptavidin conjugated to Daylight 550 (Streptavidin Protein, DyLight™ 550, Invitrogen, #84542), washed three times with 1xPBS and mounted using Fluoromount-G™ (Invitrogen, #00-4958-02) mounting medium. For pulldown experiments, genomic DNA was purified from cells using Tail buffer (10 mM Tris, 100 mM NaCl, 50 mM EDTA, 1% SDS, pH 7.5) and proteinase K digestion, followed by phenol/chloroform/isoamyl alcohol extraction. The purified DNA was fragmented by sonication using a Bioruptor Pico (Diagenode) with two cycles of 30 s on/30 s off to obtain fragments of approximately 200–500 bp. The bTMP-DNA complexes in TE buffer were immunoprecipitated overnight at 4°C using streptavidin magnetic beads (New England Biolabs, #S1420S). Beads were washed sequentially for 5 min each at room temperature with first lysis buffer (10 mM Tris-HCl, 10 mM NaCl, 1 mM EDTA, 0.2% Triton X-100, pH 8.0), wash with second lysis buffer (10 mM Tris-HCl, 150 mM NaCl, 1 mM EDTA, 0.3% SDS, pH 8.0), and TE buffer. To extract DNA and release psoralen adducts, the samples were boiled for 10 min at 90°C in 50 µl of 95% formamide containing 10 mM EDTA. The reaction volume was adjusted to 200 µl with water, and DNA was purified using the Monarch PCR and DNA Cleanup Kit (New England Biolabs, #T1030S). The resulting DNA was used as a template for qPCR.

### γH2AX foci quantification

γH2AX foci were quantified manually using Fiji software (22) after blinding the experimental groups.

### c-Fos and pCREB quantification

c-Fos and pCREB expression were quantified using Fiji software (22) after blinding the experimental groups. Quantification of pCREB-positive cells was performed using the Cell Counter plugin available within the Fiji software (22).

### Image acquisition

The images were acquired on ZEISS LSM 880 confocal microscope or Olympus AX70 microscope.

### Statistics

All statistical analyses were conducted using GraphPad Prism version 7. A p-value less than 0.05 was regarded as statistically significant.

## Discussion

TDP-43 plays a role in the repair of DNA double-strand breaks (DSBs), while its abnormal forms can induce DNA damage (6, 7). However, there is a lack of detailed knowledge regarding how TDP-43 dependent DNA damage impacts neuronal function in ALS/FTD. In this study, we have, for the first time, mapped the exact genomic locations of DSBs in neurons expressing the ALS/FTD TDP-43 associated mutants A90V and A315T, and wild-type TDP-43. This analysis revealed that a significant proportion of the affected genes by DSBs is involved in postsynaptic density organization and excitatory neurotransmission, the latest being a known contributor to ALS/FTD pathogenesis (9). Hence, the findings provide a direct link between DNA damage and aberrant neuronal transmission in ALS/FTD associated with TDP-43 pathology. Furthermore, we identified the underlying mechanism responsible for the generation of these breaks, which involves topoisomerase II β (Top2β). Our study reveals that ALS/FTD-associated TDP-43 mutants impair Top2β function, along with the neuronal capacity for activation, likely leading to extensive DNA damage.

Our study revealed that ALS/FTD-associated TDP-43 mutations A90V and A315T render neurons significantly more vulnerable to DNA breaks compared to wild-type TDP-43 on the level of actual DNA breaks. This is consistent with previous findings, assessing the general magnitude of DNA damage at the level of protein markers such as γH2AX and 53BP1 in ALS/FTD (6, 7). Notably, we found that different TDP-43 mutants A315T and A90V exert distinct effects on DSB generation, with the A90V mutation causing substantially greater genomic fragility. Interestingly, many of the genes targeted by DSBs are implicated in cellular processes previously associated with neurodegeneration such as altered neuronal transmission (9), dysregulation of the cytoskeleton (26), disrupted metabolism (27) though they had not been previously linked to DNA damage. Thus, our study revealed a novel DNA damage-dependent mechanism that may underlie the dysfunction of these processes.

Within the identified DSB gene targets we discovered those that are uniquely susceptible to DSBs in neurons expressing either the A90V or A315T TDP-43 mutants but not in wild-type TDP-43 neurons and 84 common gene targets for A90V and A315T neurons that were not present in wild-type TDP-43 samples. Among them *Grm7*, encoding metabotropic glutamate receptor 7 (mGluR7), emerged as a prominently affected gene in both mouse primary neurons and the TDP-43 rNLS mouse model. Excitatory-inhibitory imbalance is a well-established contributor to FTD/ALS disease mechanism (9, 28), however this is the first study showing that genes responsible for the glutamate signalling are targeted by DSB due to TDP-43 pathology. Interestingly, previous studies have reported a significant reduction in NMDA receptor levels in the hippocampus in Alzheimer’s disease, raising the possibility that DSBs may contribute to this decline (29). Supporting this, functional analysis of A90V-specific DSB targets revealed a significant enrichment in genes regulating glutamatergic excitatory neurotransmission. Interestingly, our findings are in line with recent research showing that excitatory neurons in the postmortem prefrontal cortex of Alzheimer’s disease (AD) patients are enriched for somatic mosaic gene fusions that arose due to DSBs (14). Similar phenomenon was reported in CK-p25 mouse model of neurodegeneration (14).

We demonstrated that neuronal activation is impaired due to ALS/FTD-associated TDP-43 A90V mutant, as evidenced by altered expression of neuronal activation markers c-Fos and pCREB. Interestingly, the A90V mutant diminishes the magnitude of pCREB expression but increases the number of activated neurons, likely desynchronized neuronal transmission compared to wild-type TDP-43. pCREB regulates the expression of genes that require prolonged transcriptional activation (20), suggesting that the A90V TDP-43 mutant may disrupt the expression of these pCREB-dependent genes.

Further, we showed that c-Fos and pCREB expression correlates with the extent of DNA damage determined by DNA damage marker, γH2AX, providing further evidence that neuronal activity is normally coupled to DNA damage. The neuronal-activity dependent generation of DSBs has been showed before (12), however this is the first study showing that TDP-43 functions in this process and that TDP-43 ALS/FTD associated mutants interfere with it.

Interestingly, etoposide treatment, a Top2β poison, revealed that the ALS/FTD-associated TDP-43 A90V mutant impairs Top2β-dependent mechanisms, likely by promoting the formation of persistent Top2β-DNA cleavage complexes. Our findings support this notion, as overexpression of TDP2 an enzyme that removes pathologically trapped Top2β-DNA complexes facilitated neuronal activation and reduced DNA damage in neurons expressing the A90V TDP-43 mutant, compared to those without TDP2 overexpression. However, we proposed that this process is responsible only for a fraction of generated DSBs, when Top2β is already bound to DNA as our BLISS data were generated at basal condition (without etoposide treatment) and TDP2 prevented from DNA damage only partially.

Our study revealed that TDP-43 is required for the binding of Top2β to DNA, and that the absence of a functional TDP-43 variant abolishes this interaction. This disruption leads to the accumulation of supercoiled DNA, likely leading to increased risk of DSBs due to unresolved torsional stress (30, 31). Topoisomerases play a critical role in resolving such stress by cleaving positively supercoiled DNA ahead of transcription bubbles or replication forks, and negative supercoils behind them (16). Thus, it is likely that TDP-43 ALS/FTD associated mutations cause the accumulation of DSBs during transcription due to inability of Top2β to resolve transcription-induced supercoiling. In line with our findings, previous studies have shown that TDP-43 disrupts chromatin dynamics downstream of the transcription start site (TSS), within the transcribed gene body, potentially interfering with efficient transcription elongation (32). Given that actively transcribed genes in neurons are known hotspots for single and double stranded DNA breaks (14, 33), the genes targeted by DSBs that we identified may be essential for ‘real-time’ neuronal function. However, due to TDP-43 pathology in ALS/FTD they undergo damage.

In our study, supercoiled DNA was still present in A90V neurons despite the large number of identified breaks compared to wild-type TDP-43, which normally should be resolved by DNA breaks. This suggests that the observed breaks may occur locally or are not sufficient to fully resolve supercoiled DNA. Supporting this interpretation, recent studies have shown that DSBs in neurons alter the three-dimensional nuclear architecture in Alzheimer’s disease models (14).

The activity of Top2β is closely linked to neuronal activation which regulates the action of Top2 β (34). It has been showed that activation of NMDA receptors triggers calcium influx, which in turn activates the phosphatase calcineurin. Calcineurin then removes phosphate groups from Top2β at serine residues 1509 and 1511, thereby enhancing its DNA-cleavage activity and promoting DSB formation (34). Further, the dysfunction of TDP-43 is enough to trigger neuronal hyperexcitability in mouse model of ALS (8). Thus, in ALS/FTD, a pathological vicious cycle may occur where altered neuronal ability for excitation caused by TDP-43 pathology disrupts Top2β function, leading to the generation of DSBs. In turn, these DNA breaks further impair neuronal signal transmission and other neuronal functions. However, given that TDP-43 also function in the repair of DSBs (6) it remains challenging to disentangle which components of this vicious cycle represent the primary cause and which are secondary effects.

Collectively, our findings provide the first comprehensive mapping of TDP-43 mutant induced DSBs in neurons and uncover a novel disease mechanism in ALS/FTD involving impaired Top2β function. These results establish a direct link between genome instability and altered excitatory neurotransmission in ALS/FTD. By identifying DSB sites and demonstrating that DNA damage can be reversed through TDP2 overexpression, this study paves the way for targeted therapeutic interventions to restore neuronal function in ALS/FTD. Genome-editing technologies, such as CRISPR based DSB repair systems, or small molecules targeting Top2β - DNA complexes, may further offer promising strategies to maintain genome stability in ALS/FTD.

## Conclusions

This study provides the first genomic map of DSBs induced by ALS/FTD-associated TDP-43 mutations in neurons. We demonstrate that mutant forms of TDP-43, particularly A90V, impair Top2β function and disrupt neuronal activation, leading to the accumulation of activity-dependent DSBs. These DNA breaks preferentially affect genes involved in excitatory neurotransmission, thereby directly linking genome instability to altered neuronal communication in ALS/FTD. Furthermore, the partial rescue of DNA integrity and neuronal activation through TDP2 overexpression highlights a potential therapeutic avenue for mitigating TDP-43 associated genomic stress. Together, our findings reveal a previously unrecognized, DNA damage-dependent mechanism contributing to neurodegeneration associated with TDP-43 pathology and open new directions for targeted strategies aimed at restoring genome stability in ALS/FTD.

## Glossary

*Double stranded breaks (DSBs)* – the type of DNA damage characterized by breaks in both strands of the DNA

*Amyotrophic lateral sclerosis (ALS)* - a progressive neurodegenerative disease in which motor neurons in the brain and spinal cord gradually deteriorate and die

*Frontotemporal dementia (FTD)* - a group of neurodegenerative disorders characterized by progressive damage to the frontal and temporal lobes of the brain

## Declarations

### Ethics approval

All experimental procedures, including animal housing, were conducted in accordance with the guidelines of the Flinders University Animal Welfare Committee (Ethics #2931) and the National Health and Medical Research Council of Australia. All work involving primary mouse neurons adhered to the Australian Code for the Care and Use of Animals for Scientific Purposes (8th edition, 2013) and received approval from the Flinders University Animal Ethics Committee (Ethics #5924).

### Consent for publication

N/A

### Availability of data and materials

The data supporting the findings of this study are included within the manuscript or in the supplementary materials. BLISS sequencing data deposited in NCBI SRA (BioSample accessions; SAMN53759388, SAMN53759389, SAMN53759390, SAMN53759391, SAMN53759392, SAMN53759393).

### Author’s contribution

Conceptualization: AK; Formal analysis AK; Finding acquisition: AK, YLC; Investigation: AK, HS, SK; Methodology: AK; Resources: AK, YLC, MLR, MF; Supervision: AK, MLR; Writing – original draft: AK; Writing – review and editing. All authors read and approved the final manuscript for publication.

## Acknowledgement

We thank Melanie Smith for her bioinformatic support.

**Supplementary material 1.** The full lists of identified DSB gene targets with their corresponding fold change values in TDP-43 mutants A315T and A90V relative to wild-type TDP-43 neurons that are presented in Figure 1.

**Supplementary material 2**. The full list of identified DSB gene targets unique for TDP-43 mutant A315T and A90V.

**Supplementary material 3**. The list of the DSB gene targets common for neurons expressing A90V and A315T TDP-43 mutants but not present in neurons expressing wild-type TDP-43 protein.

**Supplementary materials 4.** BLISS adapters and index primers used on the project.

**Supplementary materials 5.** Primers used for qPCR on the project.

**Supplementary materials 6.** Full length immunoblots and additional replications of RADAR assays

