## Supplementary material 6 for "ALS/FTD-associated TDP-43 mutations promote fragility of genes governing excitatory neurotransmission via topoisomerase IIβ impairment"

### Slide 1
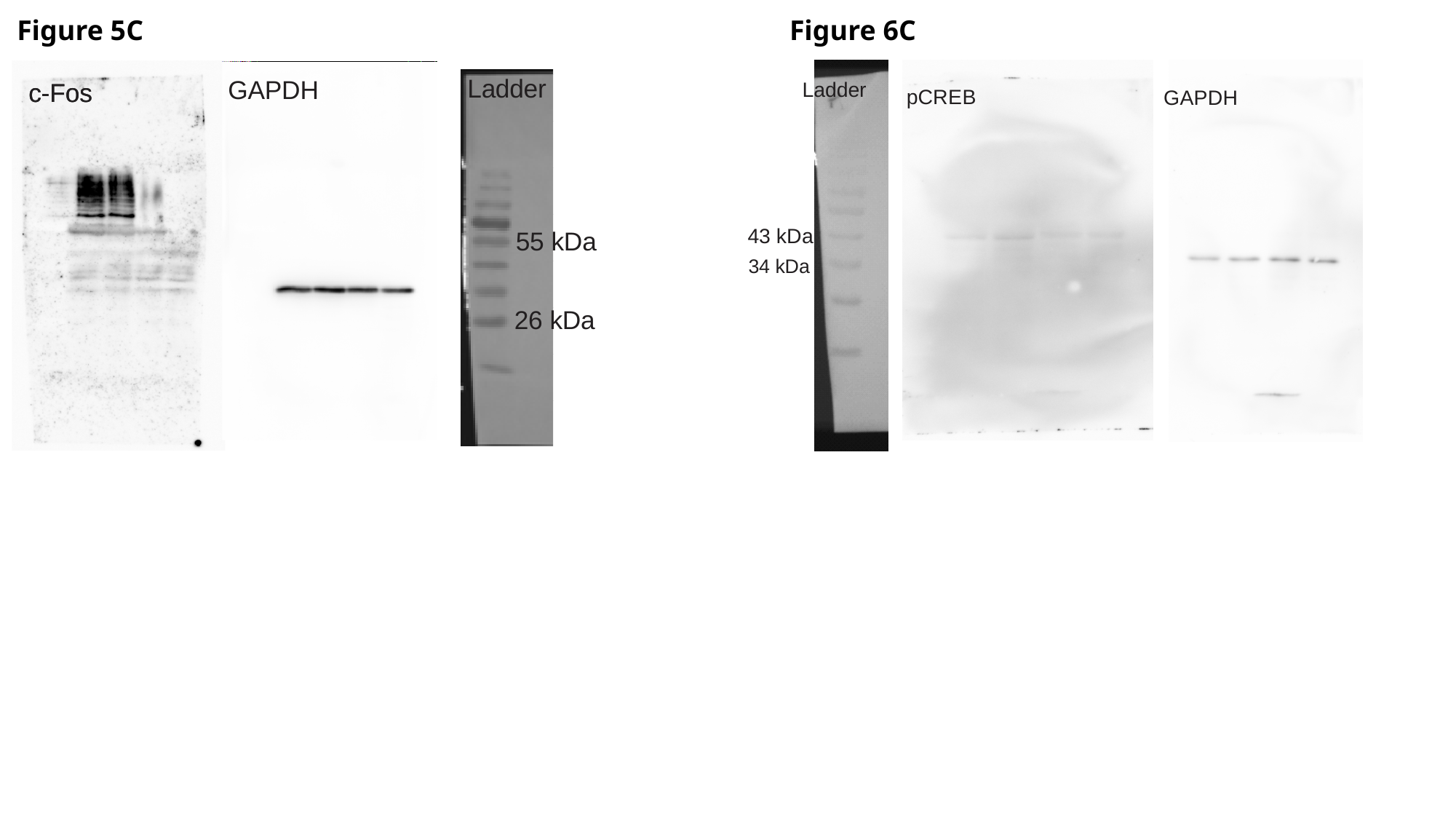

Figure 5C
Figure 6C

### Slide 2
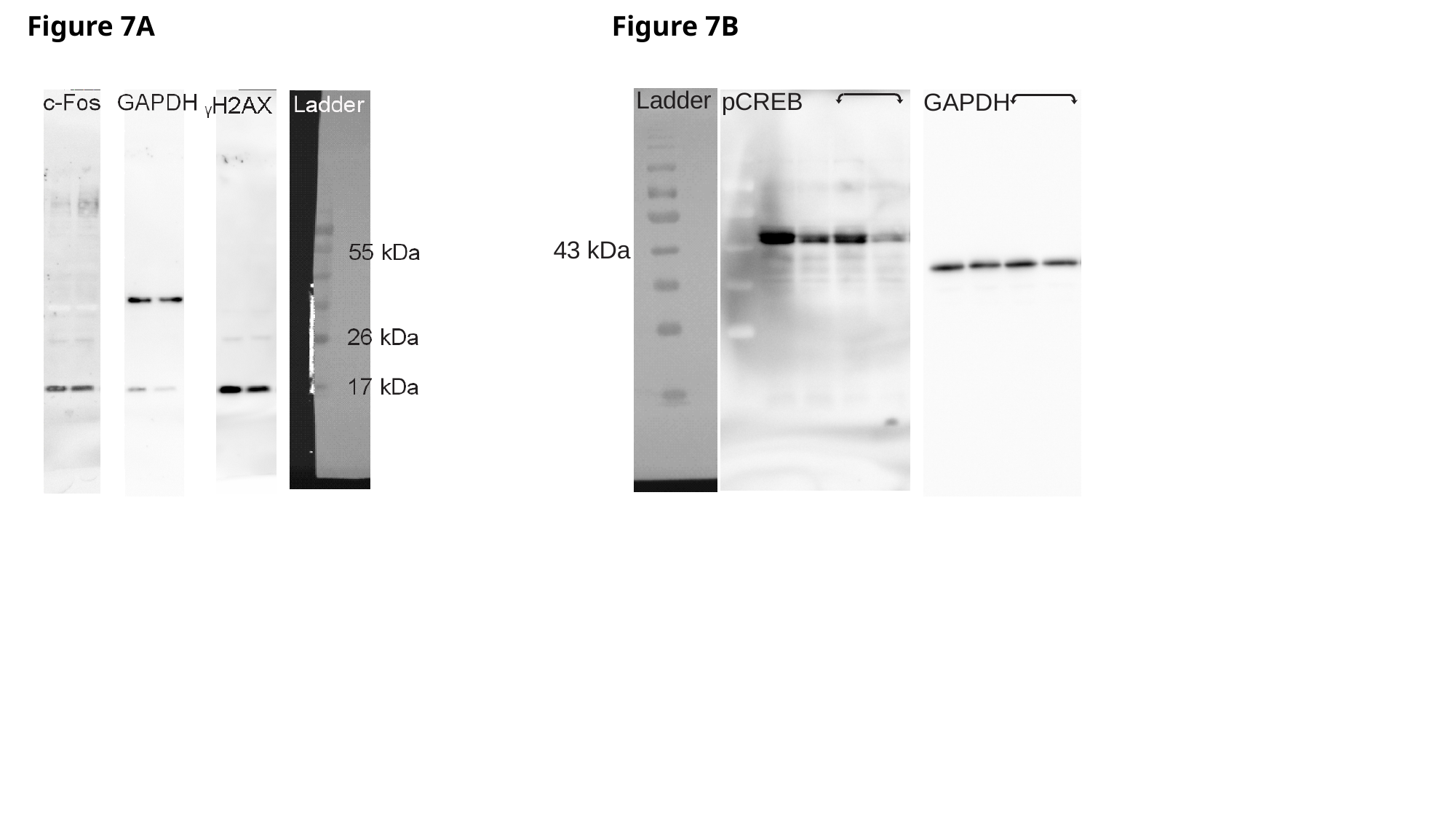

Figure 7A
Figure 7B

### Slide 3
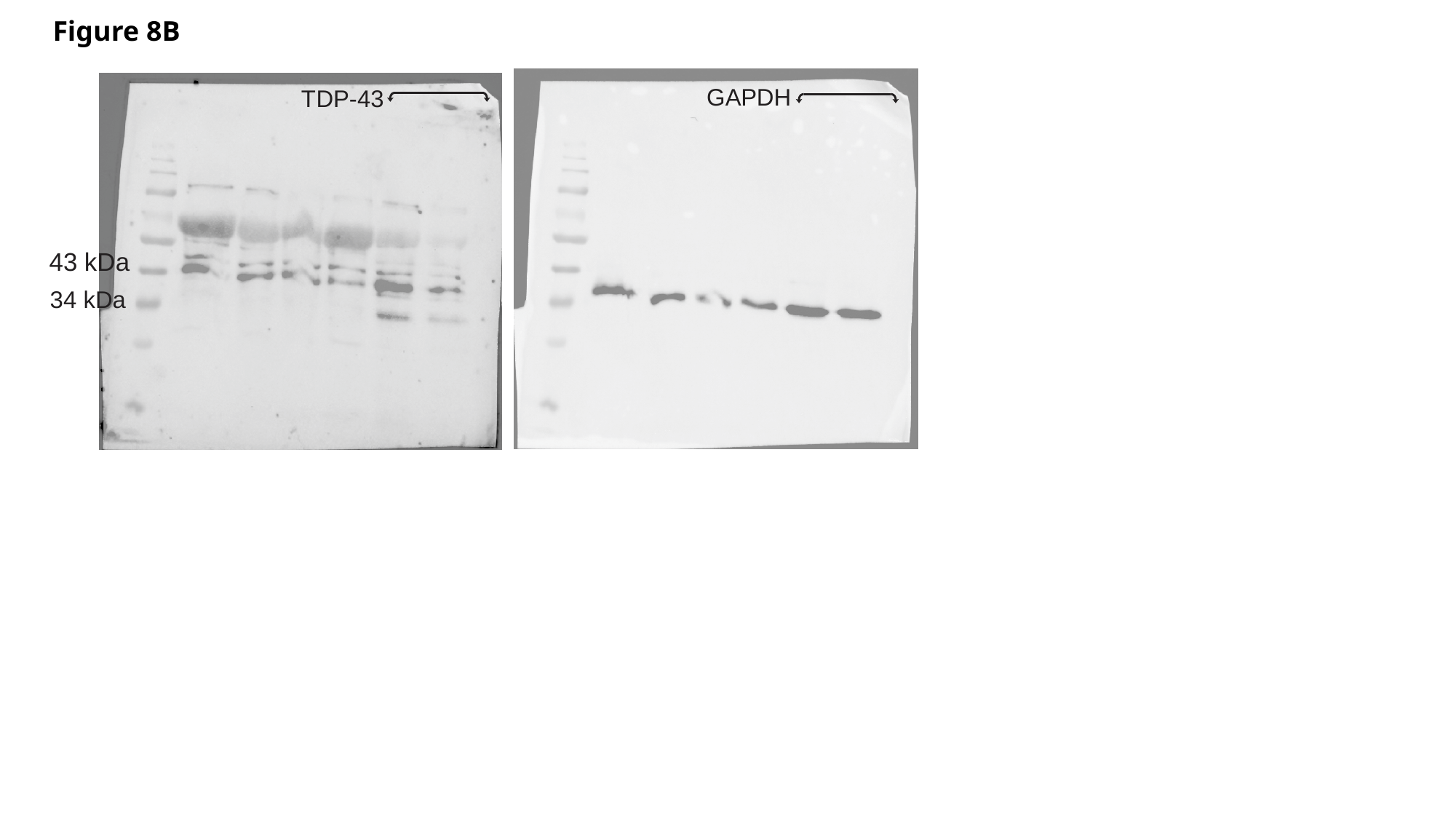

Figure 8B

### Slide 4
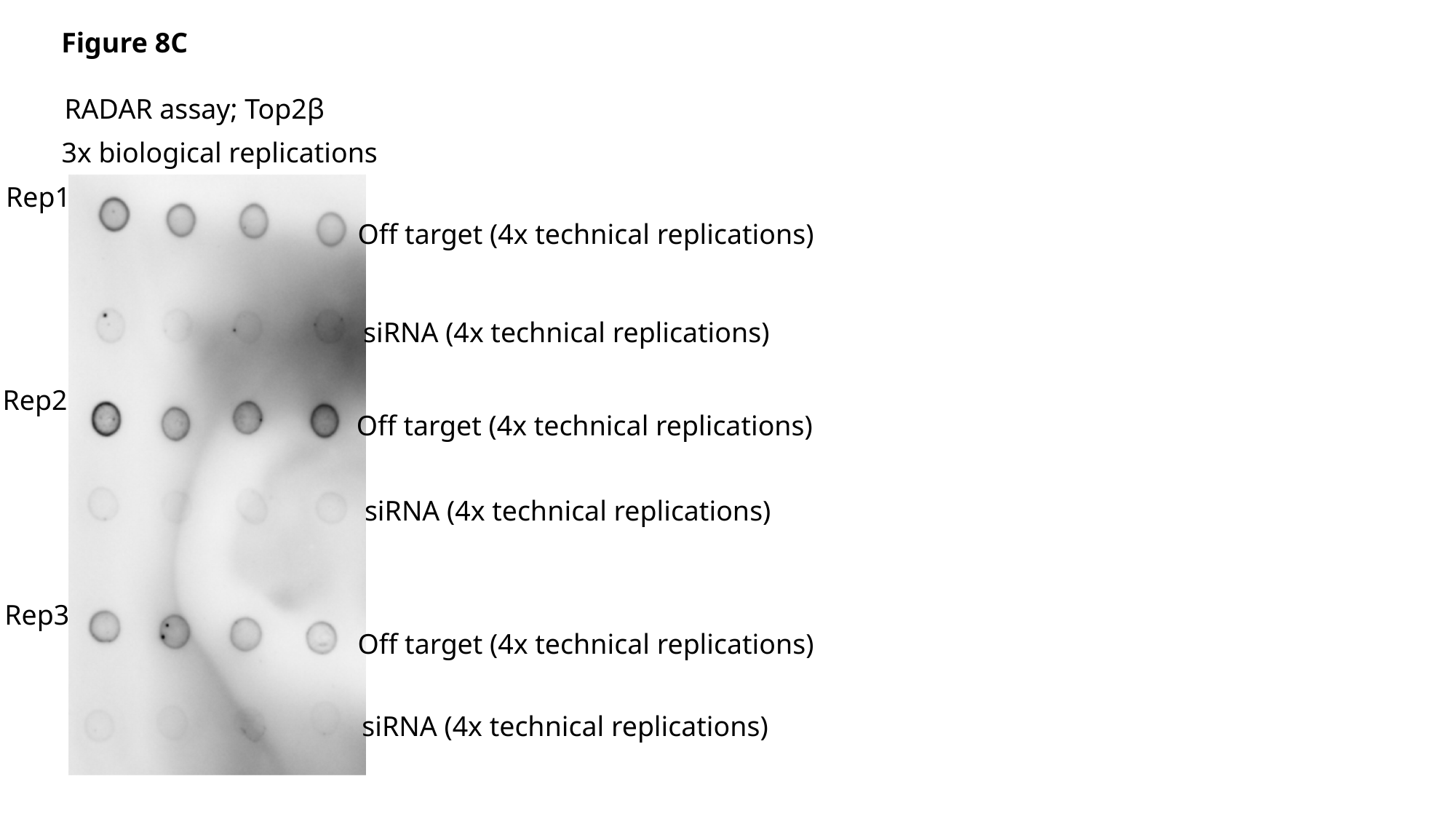

Figure 8C
RADAR assay; Top2β
3x biological replications
Rep1
Off target (4x technical replications)
siRNA (4x technical replications)
Rep2
Off target (4x technical replications)
siRNA (4x technical replications)
Rep3
Off target (4x technical replications)
siRNA (4x technical replications)

### Slide 5
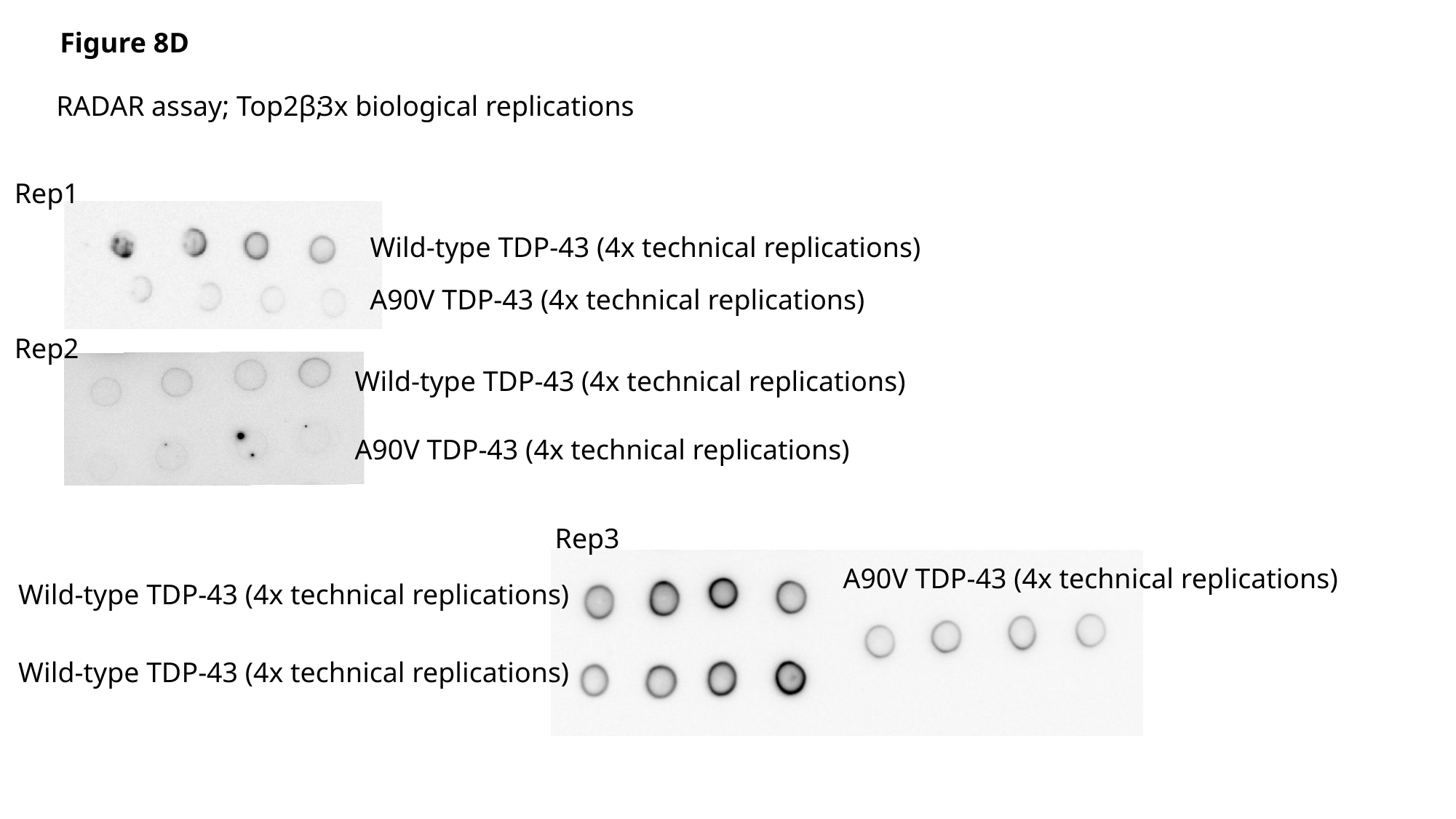

Figure 8D
RADAR assay; Top2β;
3x biological replications
Rep1
Wild-type TDP-43 (4x technical replications)
A90V TDP-43 (4x technical replications)
Rep2
Wild-type TDP-43 (4x technical replications)
A90V TDP-43 (4x technical replications)
Rep3
A90V TDP-43 (4x technical replications)
Wild-type TDP-43 (4x technical replications)
Wild-type TDP-43 (4x technical replications)
